# Distinct dendritic Ca^2+^ spike forms produce opposing input-output transformations in rat CA3 pyramidal cells

**DOI:** 10.1101/2021.10.11.463888

**Authors:** Ádám Magó, Noémi Kis, Balázs Lükő, Judit K. Makara

**Affiliations:** Laboratory of Neuronal Signaling, Institute of Experimental Medicine, 1083 Budapest, Hungary; János Szentágothai School of Neurosciences, Semmelweis University, 1085 Budapest, Hungary

## Abstract

Proper integration of different inputs targeting the dendritic tree of CA3 pyramidal cells (CA3PCs) is critical for associative learning and recall. Dendritic Ca^2+^ spikes have been proposed to perform associative computations in other PC types, by detecting conjunctive activation of different afferent input pathways, initiating afterdepolarization (ADP) and triggering burst firing. Implementation of such operations fundamentally depends on the actual biophysical properties of dendritic Ca^2+^ spikes; yet little is known about these properties in dendrites of CA3PCs. Using dendritic patch-clamp recordings and two-photon Ca^2+^ imaging in acute slices from male rats we report that, unlike CA1PCs, distal apical trunk dendrites of CA3PCs exhibit distinct forms of dendritic Ca^2+^ spikes. Besides ADP-type global Ca^2+^ spikes, a majority of dendrites expresses a novel, fast Ca^2+^ spike type that is initiated locally without backpropagating action potentials, can recruit additional Na^+^ currents, and is compartmentalized to the activated dendritic subtree. Occurrence of the different Ca^2+^ spike types correlates with dendritic structure, indicating morpho-functional heterogeneity among CA3PCs. Importantly, ADPs and dendritically initiated spikes produce opposing somatic output: bursts versus strictly single action potentials, respectively. The uncovered variability of dendritic Ca^2+^ spikes may underlie heterogeneous input-output transformation and bursting properties of CA3PCs, and might specifically contribute to key associative and non-associative computations performed by the CA3 network.

## Introduction

Dendrites play a critical role in the integration and plasticity of synaptic inputs. Voltage-dependent ion channels and passive electrical properties of dendrites enable neurons to perform various forms of linear and nonlinear input-output transformation. In particular, cortical pyramidal cell (PC) dendrites are thought to support multiple types of regenerative dendritic spikes (d-spikes), including Na^+^ spikes (mediated by voltage-gated Na^+^ channels, VGNCs), NMDA spikes (mediated by NMDARs), and Ca^2+^ spikes (mediated by voltage-gated Ca^2+^ channels, VGCCs)(Stuart and Spruston, 2015). While Na^+^ and NMDA spikes can be generated locally in individual thin dendrites of PCs (Ariav et al., 2003; Losonczy and Magee, 2006; Makara and Magee, 2013; Nevian et al., 2007; Polsky et al., 2004), Ca^2+^ spikes are thought to represent mostly global dendritic events that are responsible for bursting (Francioni and Harnett, 2021; London and Häusser, 2005; Magee and Carruth, 1999; Stuart and Spruston, 2015; Stuyt et al., 2021; Williams and Stuart, 1999).

Studies in hippocampal CA1PCs and neocortical layer 5 PCs (L5PCs) have shown that Ca^2+^ spikes are generated in the main apical trunk efficiently upon widespread synaptic depolarization in distal (tuft) dendrites concomitant with backpropagating action potentials (bAPs), and manifest as an afterdepolarization (ADP) producing a characteristic burst of additional APs (also called complex spike burst, CSB) at the soma (Harnett et al., 2013; Larkum et al., 2009, 1999; Takahashi and Magee, 2009). These results led to a concept that Ca^2+^ spikes can act as an associative dendritic signal that translates a specific input pattern (coincident activation of proximal and distal input pathways) into a burst output (a reliable form of downstream synaptic information transfer (Lisman, 1997)), and induce synaptic plasticity (Bittner et al., 2017; Takahashi and Magee, 2009). However, the question emerges: do Ca^2+^ spikes ubiquitously serve such a canonical role in PCs, or do different PC types express Ca^2+^ spikes with different properties, allowing them to support other input-output transformations and computations?

A particularly interesting PC type to explore these questions are hippocampal CA3PCs. These neurons play a fundamental role in hippocampal associative learning and memory functions (Kesner, 2013; Marr, 1971; McNaughton and Morris, 1987; Nakazawa et al., 2002; Rolls, 2007), forming a recurrent network governed by input from the dentate gyrus via mossy fibres (MFs) and the entorhinal cortex (EC)(Witter, 2007). CA3PCs frequently produce bursts of APs both *in vivo* and *in vitro* (Ding et al., 2020; Hablitz and Johnston, 1981; Hunt et al., 2018; Kowalski et al., 2016; Mizuseki et al., 2012; Oliva et al., 2016; Raus Balind et al., 2019; Wong and Prince, 1978). Investigating CSB generation in CA3PCs, we have recently reported (Raus Balind et al., 2019) that CA3PCs are heterogeneous regarding their intrinsic CSB firing propensity, and the required synaptic input patterns for evoking CSBs can be diverse as well: while in a subset of CA3PCs associative inputs drive bursts, in other CA3PCs even inputs restricted to single dendrites can efficiently produce CSBs. However, the dendritic factors underlying this diversity remained to be explored. Little is known about the generation mechanisms and properties of dendritic Ca^2+^ spikes in CA3PCs. Early studies using blind microelectrode recordings observed putative dendritic Ca^2+^ spikes to which bursting was attributed (Nuñez and Buño, 1992; Wong et al., 1979). Later, direct patch-clamp recordings in CA3PC dendrites revealed and dissected Na^+^ and NMDA spikes (Brandalise et al., 2016; Kim et al., 2012; Makara and Magee, 2013), but Ca^2+^ spikes have not been specifically examined. Intriguingly, unlike several widely studied PC types (e.g. CA1 or L5) that typically have a long or once bifurcating main apical trunk, the primary apical trunk of CA3PCs bifurcates after a relatively short distance into multiple higher-order intermediate branches. This structure creates separate apical subtrees that may represent independent, parallel integrative units for dendritic processing, allowing interactions between the radially layered inputs: detonator-type MF synapses onto proximal trunks in str. lucidum, recurrent/associative inputs onto str. radiatum dendrites and long-range EC inputs targeting distal apical branches in str. lacunosum-moleculare. However, whether individual higher-order dendritic families express Ca^2+^ spikes with specific characteristics and function remains unknown.

Here we performed patch-clamp recordings from higher-order apical dendrites of CA3PCs (alone or simultaneously with their soma) combined with two-photon (2P) Ca^2+^ imaging in the dendritic tree. We report that these dendrites support surprisingly heterogeneous forms of Ca^2+^ spikes that differ from that found in CA1PC dendrites. Besides relatively prolonged ADP-type global Ca^2+^ spikes, a majority of CA3PC dendrites expresses a novel form of fast spike (termed dendritically initiated or DI spike) that is efficiently triggered without bAPs, is mediated by a combination of fast Ca^2+^ and Na^+^ currents, and is compartmentalized to the activated dendritic family. Finally, we show that ADP-type and DI spikes produce opposing forms of somatic output: bursts versus strictly single APs, and thereby may actively promote different firing modes in morpho-functionally different CA3PCs. The unique properties of DI spikes, such as their fast kinetics and anti-bursting effect fundamentally differ from the classical associative role of dendritic Ca^2+^ spikes, suggesting that they may enrich the computational repertoire of CA3PCs with novel, cell type- or circuit-specific functions.

## Results

### Identification of regenerative spikes in CA3PCs dendrites

To directly investigate active dendritic properties of CA3PCs we performed dual soma-dendrite current-clamp recordings combined with two-photon imaging in dendrites (**Fig. 1a**). Dendritic recordings were made from higher order trunks at distances ∼165-400 μm from the soma. Subthreshold steady-state voltage signals attenuated more strongly from dendrite to soma than from soma to dendrite, as expected (**Fig. 1b,d,** n=15, p=0.004, Wilcoxon test).

**Figure 1.**
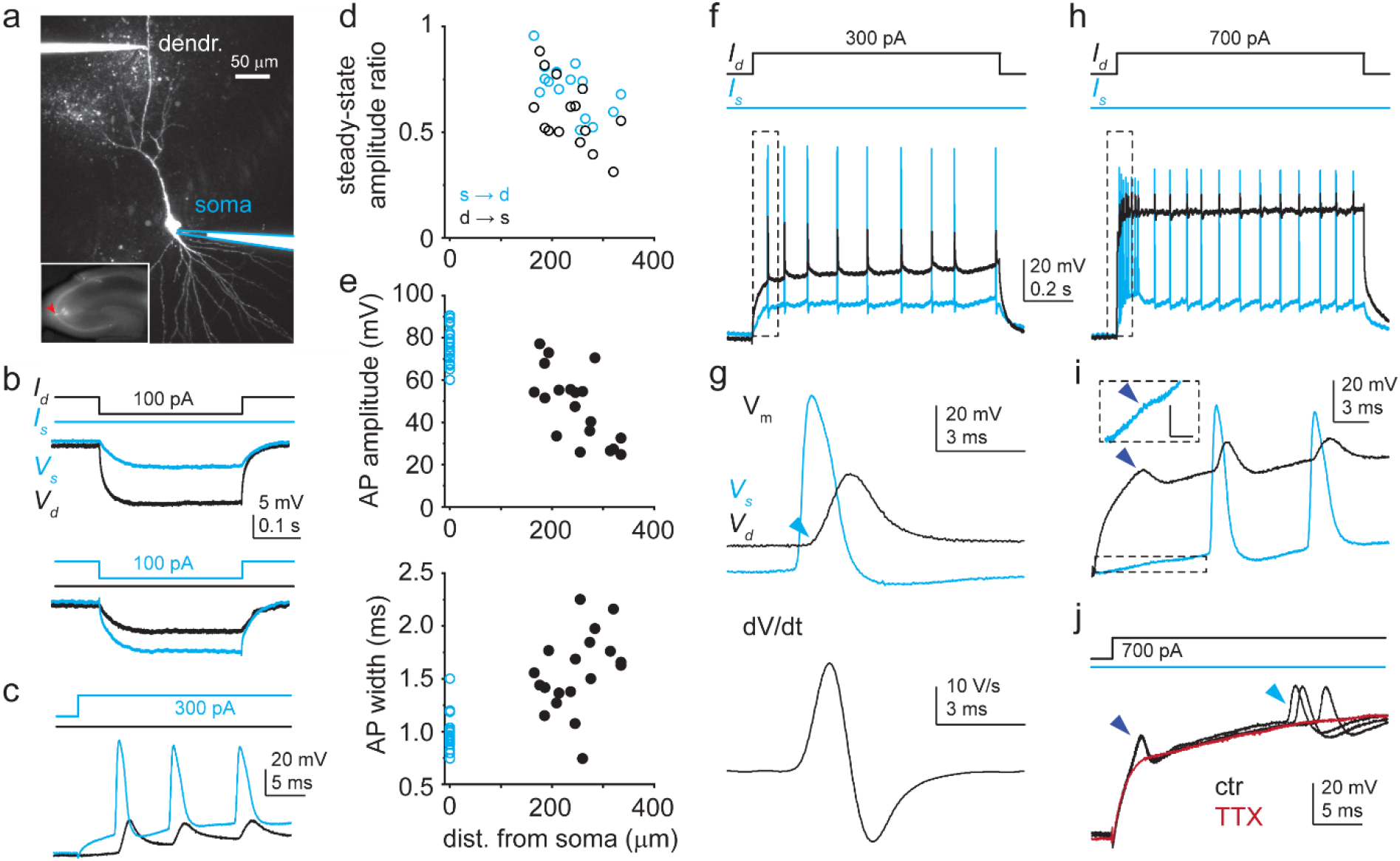
Dendritic Na^+^ spikes in dual recordings of soma and dendrite of CA3PCs. **a)** 2P collapsed z-stack image of a dually recorded CA3PC (dendritic pipette distance: 255 μm). Inset: soma (red arrowhead) located in CA3. **b)** Somatic (blue) and dendritic (black) voltage responses (V_s_ and V_d_) to negative step I_inj_ into the dendrite (top) and the soma (bottom). I_inj_ protocols shown on the top. **c)** Positive I_inj_ at the soma evokes APs (blue) that backpropagate to the dendrite (black). **d)** Voltage transfer between soma and dendrite (n=15 experiments). **e)** Dendritic distance dependence of amplitude and width at half amplitude of the first bAP evoked by somatic I_inj_ (n=19 experiments). **f)** Voltage responses at the dendrite (black) and at the soma (blue) to a 1-sec-long 300 pA dendritic I_inj_ in the cell shown in a. **g)** First regenerative event in f (box) is a bAP. Top: voltage; bottom: corresponding dendritic dV/dt. Note the sudden start of the bAP (“kink”, indicated by light blue arrowhead). **h)** Dendritic and somatic voltage responses to 1-sec-long 700 pA dendritic I_inj_ in the cell shown in a. **i)** Initial part of the voltage response in h (box) enlarged. A short-latency regenerative dendritic event (dark blue arrowhead) precedes bAPs. Attenuated somatic response is shown enlarged in a dashed box. Scale bar in box: 5 mV, 3 ms. Similar events were observed in 3 other CA3PCs. **j)** Short-latency dendritic spikes and APs (black traces; 3 repetitions) are eliminated by 1 μM TTX (dark red). **Fig. 1 – figure supplement 1. Characteristics of different regenerative dendritic events**

Applying 1-sec-long depolarizing current injections via either the dendritic or somatic electrode, we readily identified two types of previously described fast VGNC-mediated regenerative voltage responses in the dendrites: bAPs and dendritic Na^+^ spikes (Kim et al., 2012). Backpropagating APs could be observed in almost all dendrites both by sufficient somatic or dendritic depolarizing stimuli; they followed somatic APs with short latency, their amplitude decreased gradually with distance after the first ∼150 μm (Kim et al., 2012), and they displayed a canonical sharp initiation profile („kink”, Gidon et al., 2020; Smith et al., 2013) and short duration, all characteristic for bAPs (**Fig. 1** and **Fig. 1 - figure supplement 1)**. On the other hand, fast dendritic Na^+^ spikes were generated in a subset of experiments (n=4) by large (I_inj_; ≥600 pA) current injections into the dendrite. These events appeared with very short latency (<7 ms) on the initial depolarizing phase of the step, and attenuated strongly from dendrite to the soma (**Fig. 1h-j**), as previously described (Kim et al., 2012). Both of these fast spike types disappeared after the application of 1 μM TTX in the bath, confirming that they were mediated by VGNCs (**Fig. 1j**, n=12 experiments; local spikes were present in 2 of these experiments under control conditions; see also later the effect of TTX in single-site dendritic recordings).

In the majority of dual recordings (17 of 21) we also observed slower regenerative voltage responses that we considered to be putative dendritic Ca^2+^ spikes (**Fig. 2**). These spikes were elicited with relatively longer latency or at the steady-state depolarized phase, and had highly heterogeneous kinetics (see below). We classified putative Ca^2+^ spikes into two broad groups based on their initiation characteristics. The first, “classical” group consisted of regenerative ADP forms, i.e. responses that followed a bAP with an additional voltage peak (**Fig. 2b-d**). ADPs had a wide range of kinetics. In some dendrites the ADP manifested as a sustained afterdepolarization that triggered additional APs and gradually built up a prolonged, slowly decaying depolarization driving a burst of APs („slow ADP”, >∼50-100 ms, **Fig. 2b**). In other cases, the ADP occurred as a transient regenerative voltage response following 1-3 APs („fast ADP”, **Fig. 2c-d**), which either remained subthreshold or evoked additional AP(s) at the soma.

**Figure 2.**
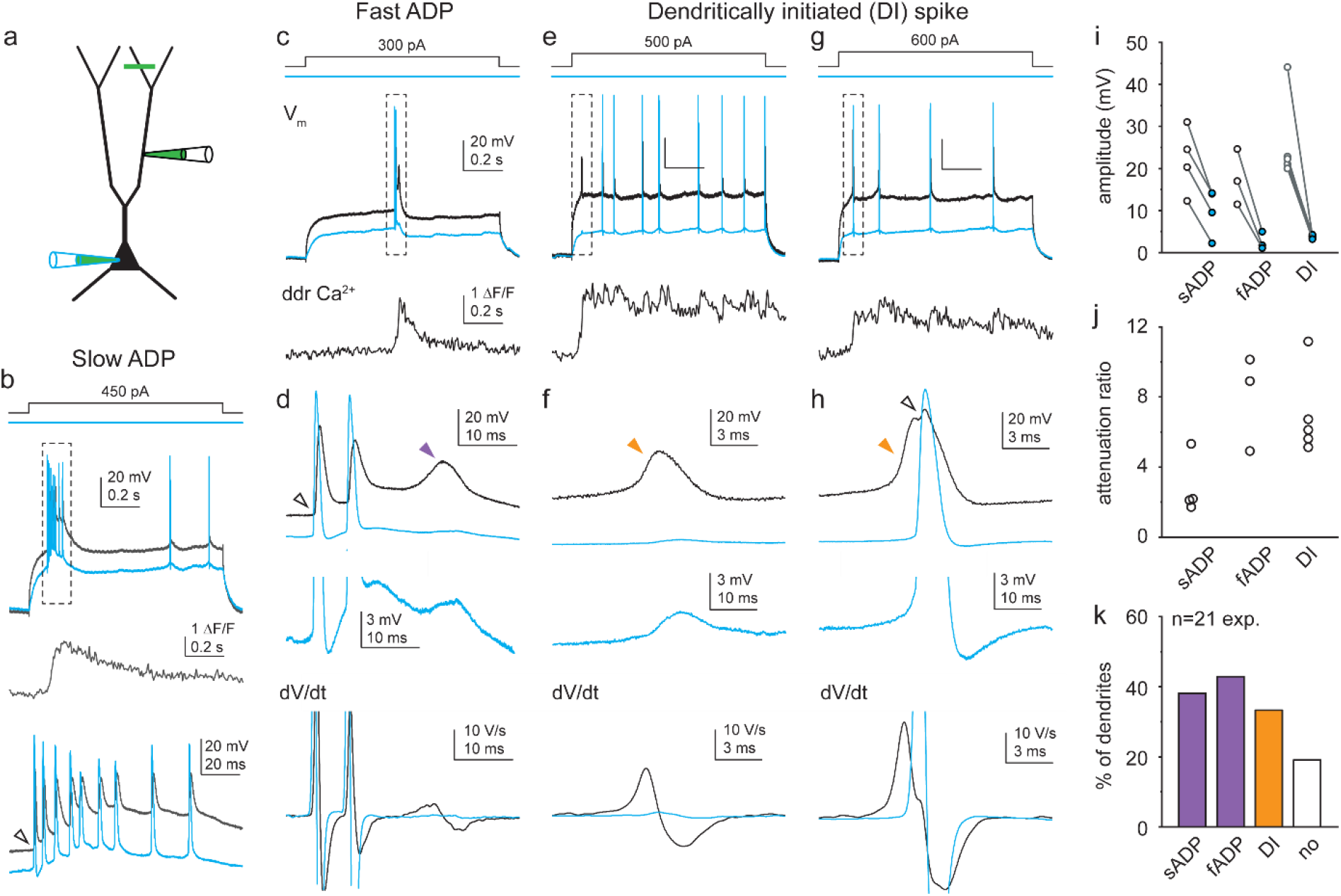
Diverse dendritic Ca^2+^ spike forms in soma-dendrite dual recordings from CA3PCs. **a)** Schematic of experiments. Green line indicates typical Ca^2+^ imaging site distal from the dendritic patch. **b)** Representative recording of a slow ADP. Dendritic (black) and somatic (blue) voltage response pair to dendritic I_inj_ (top), and corresponding distal dendritic Ca^2+^ signal (middle). Dashed box on top indicates the event that is enlarged on the bottom. Note the prolonged sustained ADP building a slow depolarization that is larger in the dendrite. Open arrowhead denotes kink of the initiating bAP. **c)** Representative recording of a fast ADP. **d)** Event in dashed box in c is shown enlarged. Top, dendritic (black) and somatic (blue) voltage pair. Middle, somatic trace magnified. Bottom, corresponding dV/dt traces (APs truncated). Open arrowhead denotes “kink”; purple arrowhead points to the fast ADP. **e-g)** Representative recordings of dendritically initiated (DI) spikes, either isolated in the dendrite (**e-f**) or evoking a consecutive AP (**g-h**). Panels as in c-d. Open arrowhead denotes AP with “kink”; orange arrowheads indicate DI spikes. Traces in b,c,e,g are from different cells. Note the different time scales of various spike forms. **i)** Ca^2+^ spike amplitudes in dendrite (open circle) and soma (blue circle) in individual recordings. See Materials and Methods for details. **j)** Calculated attenuation (Ampl_ddr_/Ampl_soma_) in individual recordings. **k)** Propensity of different Ca^2+^ spike forms (in total n=21 dendrites with dual recordings).

In addition to “classical” ADPs, we discovered a second, unconventional form of putative Ca^2+^ spikes. These events were generated without an initiating bAP, and were therefore termed dendritically initiated (DI) spikes. The rise of these spikes typically followed a concave, often exponentially developing profile that could be well distinguished from the sharp „kink” of bAPs (**Fig. 1 – figure supplement 1b-e**). Although DI spikes did not require bAPs for their initiation and could be evoked in isolation in the dendrite (**Fig. 2e-f**), they could also evoke consecutive bAPs (**Fig. 2g-h**). The dendritic origin of both ADPs and DI spikes was confirmed by their strong attenuation towards the soma (**Fig. 2i-j**, dendrite-soma attenuation ratio: ADPs: 5.06 ± 1.28, n=7; DI spikes: 6.97 ± 1.09, n=5). ADPs and DI spikes co-occurred in some of the recordings (**Fig. 2k**).

Simultaneously with electrophysiology, we measured dendritic Ca^2+^ signals (using OGB-1 or OGB-6F, see Materials and Methods) at locations >89 μm (mean ± SEM.: 234 ± 20 μm) distal from the dendritic pipette. All forms of the above identified putative Ca^2+^ spikes (but not local Na^+^ spikes, **Fig. 1 - figure supplement 1a**) were accompanied by large distal dendritic Ca^2+^ signals that coincided with the onset of the voltage response (**Fig. 2b,c,e,g**), suggesting that the voltage signals propagated actively and involved regenerative Ca^2+^ influx.

### Characterization of dendritic Ca^2+^ spikes

Equipped with the fingerprint characteristics to identify putative Ca^2+^ spikes (see also **Fig. 1 – figure supplement 1**), we further characterized the properties of these spikes using I_inj_ in a larger set of single-site dendritic recordings (n=69, at 159-450 μm from soma, mean ± SEM: 272 ± 8 μm). Replicating the findings of dual recordings, 1-second-long dendritic I_inj_ steps elicited different types of putative Ca^2+^ spikes, i.e. ADPs and/or DI spikes in the majority (∼85%) of individual dendrites, in addition to bAPs and dendritic Na^+^ spikes (**Fig. 3a-d, f**, and **Fig. 1 – figure supplement 1**). Putative Ca^2+^ spikes were often generated repetitively, and in some cells their kinetics were variable even across repetitions (**Fig. 3 – figure supplement 1**). The spikes were unaffected by blockade of AMPA and NMDA receptors, confirming that they were not related to excitatory synaptic activity (**Fig. 3 – figure supplement 2**).

**Figure 3.**
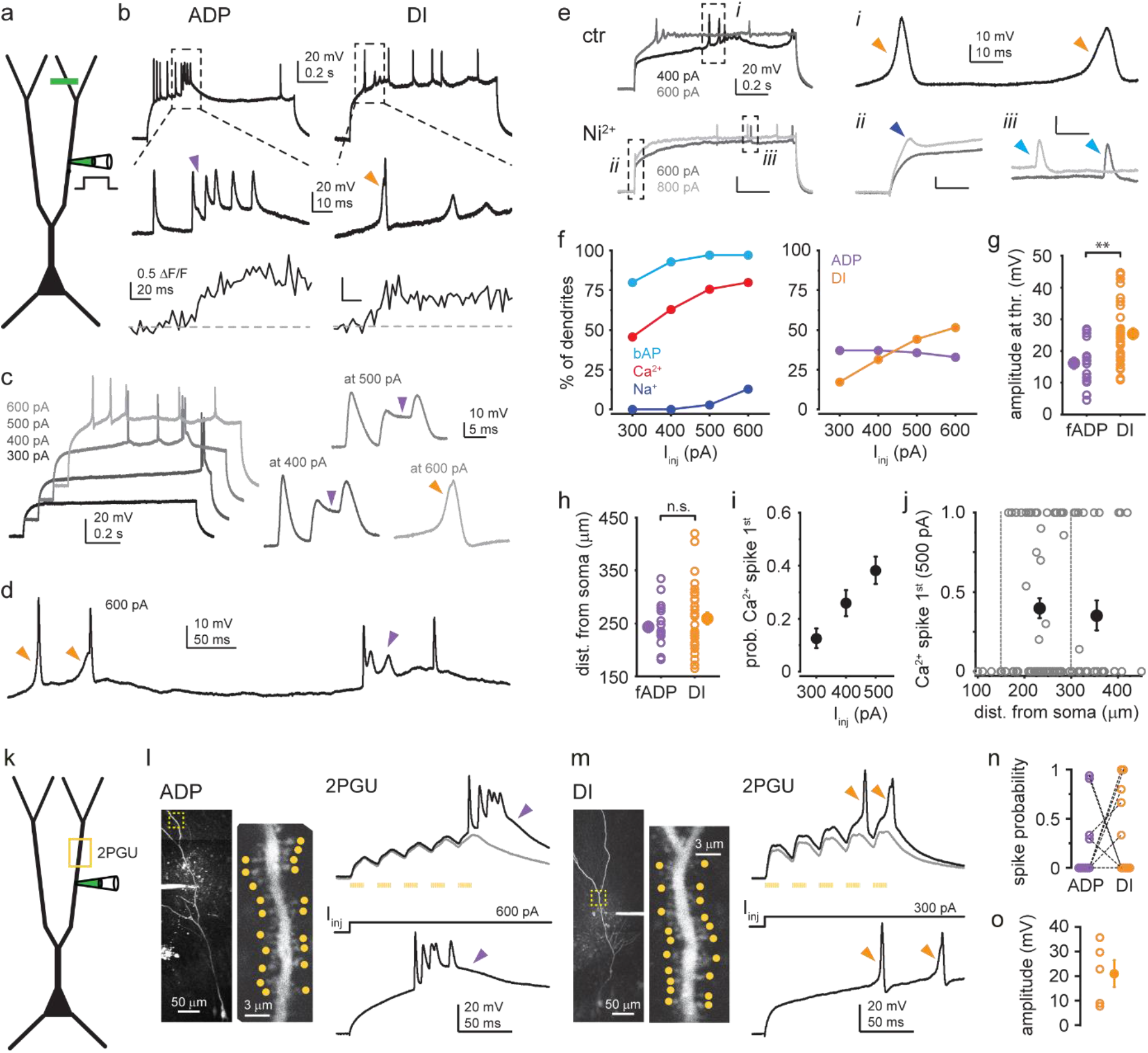
Characterization of dendritic Ca^2+^ spike types. **a)** Schematic of dendrite-only experiments. **b)** Representative dendritic voltage and Ca^2+^ recording of ADP (left) and DI spike (right) from two different cells. **c)** Example responses of a dendrite to increasing I_inj_. Right: Ca^2+^ spikes expressed on different traces magnified. Purple arrowheads: ADP; orange arrowhead: DI spike. **d)** Single trace in response to 600 pA I_inj_, containing heterogeneous Ca^2+^ spike forms. **e)** Representative traces under control conditions (top, two different I_inj_) and after bath application of 200 μM Ni^2+^ (bottom). Dashed boxes are enlarged on the right to show DI spikes (i, orange arrowhead), dendritic Na^+^ spike (ii, dark blue arrowhead) and bAPs (iii, light blue arrowhead). Note that Na^+^ spikes were resistant to Ni^2+^ (see also Fig. 1 – figure supplement 1i). **f-j)** Summary of Ca^2+^ spike properties (dendrite-only and dual recordings pooled). **f)** Left, percent of dendrites expressing bAPs, Ca^2+^ spikes (ADPs and DI spikes included) and Na^+^ spikes to 300-600 pA I_inj_ (n=70 dendrites). Right, percent of dendrites expressing different types of Ca^2+^ spikes (ADPs and DI Ca^2+^ spikes) to 300-600 pA I_inj_ (n=70 dendrites). **g)** Amplitude of fast ADPs and DI spikes at threshold I_inj_. Open circles: mean amplitude in individual dendrites; filled symbol: mean ± SEM of experiments. **h)** Dendritic distance of pipette from the soma in the experiments in g. **i)** Mean probability (range: 0-1) that Ca^2+^ spike was the first regenerative event evoked by 300, 400 or 500 pA I_inj_ (n=70, 75, 82 dendrites). **j)** Probability (range: 0-1) that Ca^2+^ spike was the first regenerative event evoked by 500 pA I_inj_, as a function of pipette distance from soma. Open grey circles: individual dendrites; filled black symbols: mean ± SEM of measurements in 151-300 μm (n=56) and 301-450 μm (n=26) distance range from soma. **k)** Schematic of 2PGU experiments. **l)** Left, z-stack of a CA3PC, and single scan of the dendritic segment (marked by yellow dashed box) indicating the 20 synapses stimulated by 2PGU (yellow dots). Right, top: example responses to 2PGU (20 spines stimulated quasi-synchronously 5x@40 Hz). Grey: subthreshold, black: suprathreshold response. Bottom: same dendrite responding to I_inj_ via the pipette. **m)** Same as **l** for a dendrite with DI spike. Note the similarities in spike types by 2PGU and I_inj_ in l-m. **n)** Relative probability of evoking ADPs and DI spikes in dendrites (number of traces displaying the respective d-spike divided by the total number of suprathreshold traces). Dashed lines connect data from the same dendrites. **o)** Amplitude of DI spikes evoked by 2PGU was similar to that typical with I_inj_ (compare to panel g). **Fig. 3 – figure supplement 1. Variability of Ca^2+^ spike phenotypes across and within CA3PCs** **Fig. 3 – figure supplement 2. Dendritic Ca^2+^ spikes do not depend on excitatory synaptic activity** **Fig. 3 – figure supplement 3. Dendritic Ca^2+^ spikes in CA1PCs are ADP-type**

In a set of dendrites we systematically determined the propensity of the two main Ca^2+^ spike types evoked in the 300-600 pA I_inj_ range and we found that ADPs and DI spikes were expressed in 56 and 53 % of the investigated dendrites, respectively (**Fig. 3f**, n=70 cells, dual- and single-site experiments pooled), and in 24 % they both were present. The dominant Ca^2+^ spike type often depended on the I_inj_ level, with ADPs evoked at smaller and DI spikes evoked at higher I_inj_ (**Fig. 3c, f**). Furthermore, in some dendrites different Ca^2+^ spike types occurred intermingled within the same depolarizing trace (**Fig. 3d**). As in dual recordings, both ADPs and DI spikes were associated with large Ca^2+^ signals measured in dendritic segments >80 μm (mean ± SEM: 215 ± 9 μm) distal to the patch pipette (**Fig. 3b**). Bath application of 200 μM

Ni^2+^ eliminated all types of putative Ca^2+^ spikes, whereas bAPs and dendritic Na^+^ spikes were spared (**Fig. 3e**, and **Fig. 1 – figure supplement 1i**). These results confirmed that VGCCs played a fundamental role in mediating ADPs and DI spikes. Interestingly however, the amplitude of DI spikes, measured at threshold I_inj_ (see Materials and Methods for amplitude measurement criteria), was larger (25.4 ± 1.7 mV, n=30) than that of fast ADPs (16.2 ± 1.8 mV, n=18; Mann-Whitney test p=0.003, **Fig. 3g**). This difference was not explained by different dendritic distance of the recordings from the soma (DI spikes: 259 ± 12 μm, ADPs: 243 ± 10 μm, Mann-Whitney test p=0.601, **Fig. 3h**), and the amplitude of neither spike type correlated with distance (Spearman correlation: ADP: R=-0.260, p=0.296, n=18; DI: R=-0.297, p=0.110, n=30), suggesting a difference in the spike generation mechanism rather than simple distance-dependent variation of spike properties.

We next addressed the question of whether the soma or the dendrite is more likely to first generate a regenerative spike upon dendritic depolarization. When the patch pipette was positioned at relatively proximal dendritic locations (<∼150 μm from soma), somatic APs were always evoked first by dendritic depolarization. However, at more distal dendritic locations (>∼150 μm from soma), DI Ca^2+^ spikes were often evoked before bAPs, and this propensity depended on the strength of I_inj_ (**Fig. 3i**, ratio at 300 pA: 0.13 ± 0.04, n=70; 400 pA: 0.26 ± 0.05, n=75; 500 pA: 0.38 ± 0.05, n=82; p=0.001, Kruskal-Wallis test) but was independent from the distance from soma (**Fig. 3j**, Spearman correlation R=-0.031, p=0.778, n=82). Thus, DI spikes can be evoked efficiently by local depolarization of medial-distal apical trunks of CA3PCs without any preceding somatic activity.

To test whether the distinct forms of Ca^2+^ spikes observed by I_inj_ could be also evoked by more physiological forms of stimulation, we next performed experiments employing synaptic stimulation on the trunk using 2P glutamate uncaging (2PGU; **Fig. 3k-l**). We patched dendrites (n=10 experiments, 230 ± 17 μm from soma) and stimulated 20 clustered spines located along the same dendrite at 171-441 μm distance from the soma quasi-simultaneously 5x at 40 Hz by adjusting the laser power to produce moderately suprathreshold stimulation (bAP or dendritic spike evoked by any of the last three of the five stimuli). To avoid activation of confounding slow NMDA spikes, these experiments were performed in the presence of an NMDAR blocker in the bath (D-AP5, 50 μM). Synaptic stimulation was able to elicit characteristic and well distinguishable ADPs and DI spikes (**Fig. 3n**) either separately (ADP only: 3/10 dendrites, **Fig. 3l**; DI spike only: 4/10 dendrites, **Fig 3m**) or in combination (1/10 dendrites). In the two dendrites with no d-spikes by moderate stimulation, stronger stimuli could activate ADPs (data not shown). The Ca^2+^ spike profile of the dendrite evoked by uncaging was similar to that seen with I_inj_ via the pipette (**Fig. 3l-m**), and DI spikes had amplitudes comparable to that typically evoked by I_inj_ (**Fig. 3o**). These results confirm that the Ca^2+^ spike form is an inherent property of the dendrite, and separate Ca^2+^ spike modes can be elicited by wide range of stimuli that reach sufficiently strong local depolarization.

The surprisingly variable properties of putative dendritic Ca^2+^ spikes found in CA3PC dendrites prompted us to compare these spikes to the well-characterized Ca^2+^ spikes of CA1PCs (Golding et al., 1999; Magee and Carruth, 1999; Takahashi and Magee, 2009). Therefore, we performed similar experiments in the apical trunks of CA1PCs, at 227-457 μm distance from soma (n=12, **Fig. 3 – figure supplement 3**). In most CA1PCs, dendritic depolarization by 300-600 pA I_inj_ evoked VGCC-mediated ADPs (**Fig. 3 – figure supplement 3a-c, g**), but the required I_inj_ to trigger the spikes was higher than that in CA3PCs (**Fig. 3 – figure supplement 3d**). Furthermore, DI spikes were not observed in CA1PC trunks (**Fig. 3 – figure supplement 3e**), and accordingly, bAPs were always evoked before Ca^2+^ spikes (**Fig. 3 – figure supplement 3f**).

Thus, active properties of CA3PC trunk dendrites fundamentally differ from those of CA1PCs, indicating that the repertoire and roles of dendritic computation can vary in different types of rat PCs. Curiously however, some features of CA3PC dendritic Ca^2+^ spikes resembled those recently described in L2/3PCs in cortical slices removed from human patients (hL2/3PCs; Gidon et al., 2020). Specifically, the DI spikes we observed in CA3PCs shared several properties with a novel type of Ca^2+^ spikes in hL2/3 PCs: they were characterized by bAP-independent initiation, fast rise and large amplitude (**Fig. 4a-b**), which inversely scaled with increasing I_inj_ (**Fig. 4c-e**). In some dendrites (6 out of 12 tested), the depolarization level-dependent reduction in amplitude created a window of dendritic stimulus strength where APs could be specifically triggered by these dendritic events (**Fig. 4f**), similarly to that described in hL2/3PCs (Gidon et al., 2020). A systematic comparison of the parameters of DI spikes in CA3PCs to those reported for hL2/3PCs revealed remarkable qualitative similarities with moderate quantitative differences (CA3PC: amplitude at threshold I_inj_: 25.43 ± 1.74 mV, n=30; width at half amplitude: 6.43 ± 0.75 ms, n=18; exponential decay constant (tau) of normalized amplitude vs I_inj_: 0.47; hL2/3PC parameters from (Gidon et al., 2020): amplitude: 43.8 ± 13.8 mV; width: 4.4 ± 1.4 ms; tau: 0.39). We conclude that DI spikes are expressed in specific neuron types of various mammalian species including humans and rats.

**Figure 4.**
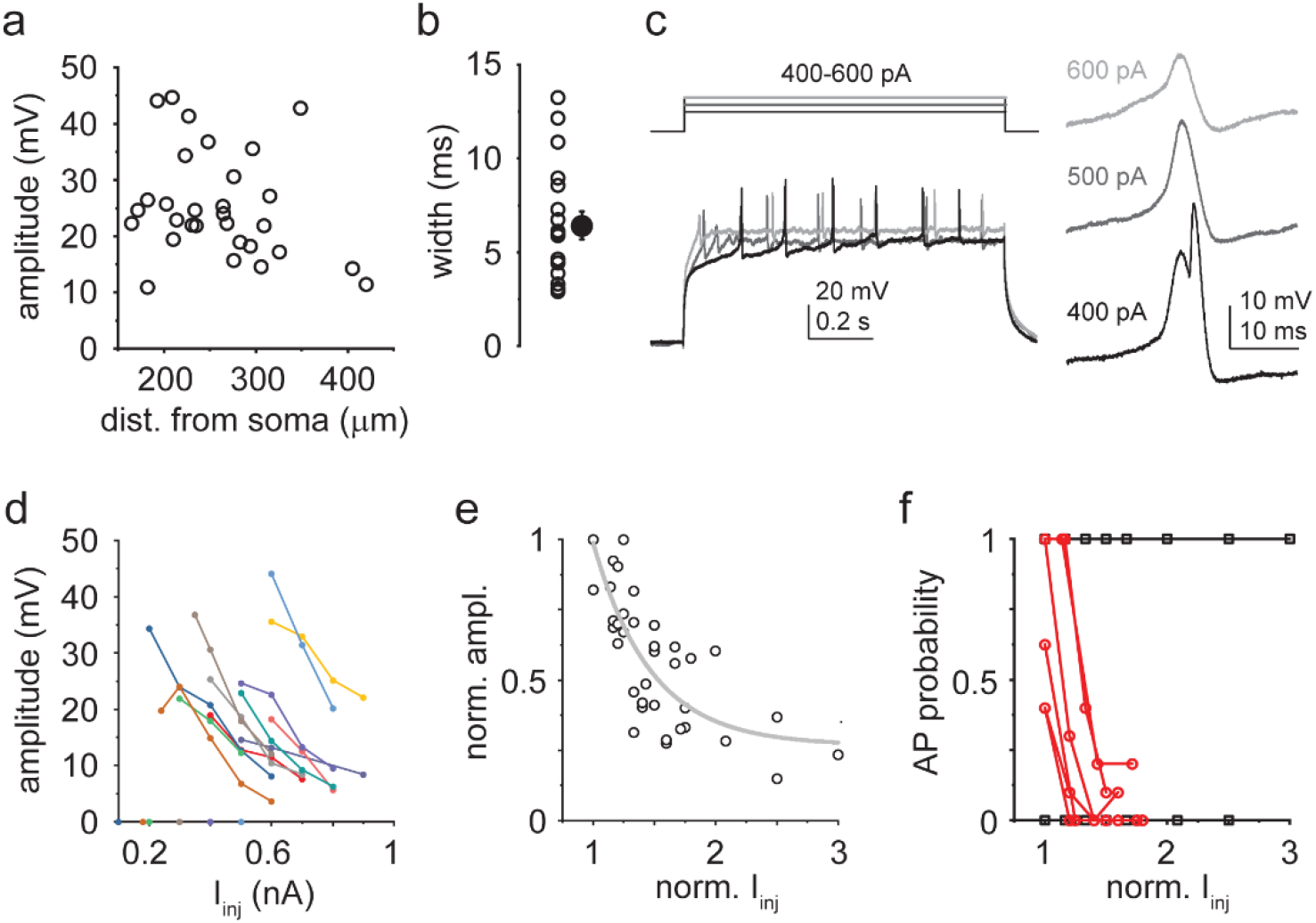
Properties of DI Ca^2+^ spikes. **a)** Spike amplitude as a function of pipette distance from soma (circles: individual dendrites, n=30). **b)** Width at half amplitude of DI spikes. Open circles: individual dendrites; filled symbol: mean ± SEM (n=18). **c)** Representative responses of a dendrite to 400-600 pA I_inj_. Right, first regenerative events enlarged. **d)** I_inj_-amplitude response curves in 12 dendrites with DI spikes. **e)** Normalized I_inj_-amplitude relationship established from the data in d. Grey line: exponential decay function. **f)** Probability of AP firing directly elicited by DI spikes at various I_inj_ levels. In 6 out of 12 dendrites, larger I_inj_ evoked DI spikes with progressively lower AP probability (red). In the other 6 dendrites AP probability remained 0 or 1.

### Ion channel mechanisms

To study the properties of CA3PC Ca^2+^ spikes in isolation, we bath applied the VGNC inhibitor TTX (1 μM). As expected, TTX completely eliminated bAPs and Na^+^ spikes (**Fig. 1 – figure supplement 1h-i**, see also **Fig. 1**), whereas putative Ca^2+^ spikes, associated with large dendritic Ca^2+^ signals, persisted. Interestingly, the kinetics of the regenerative spikes remaining in TTX varied in a wide range, but mirrored the behavior in ACSF. That is, in dendrites that under control conditions expressed only ADPs (fast or slow) but no DI spikes, TTX-resistant Ca^2+^ spikes were typically slow (**Fig. 5a, c**, width: 166.6 ± 26.8 ms, n=12), whereas those dendrites that fired DI spikes (with or without ADPs) under control conditions displayed dominantly fast, transient, TTX-resistant Ca^2+^ spikes (**Fig. 5b, c**, width: 9.3 ± 0.7 ms, n=13, Mann-Whitney test p<0.001), although a smaller slow component was often also present. In some cases (mostly in cells expressing both ADPs and DI spikes in ACSF) slow and repetitive fast components were mixed (**Fig. 5 – figure supplement 1a**). The duration of TTX-resistant Ca^2+^ spikes in CA3PCs clearly separated from those of CA1PC dendrites under similar conditions (width: 39.4 ± 5.1 ms, n=6, **Fig. 3 – figure supplement 3h-i**), further confirming cell-type specific differences in Ca^2+^ spike properties. Finally, in ∼25% of dendrites, no clear regenerative voltage responses could be evoked after TTX application (**Fig. 5 – figure supplement 1b**). These results, together with the elimination of all types of Ca^2+^ spikes by 200 μM Ni^2+^ (**Fig. 3e**) confirm a fundamental role of VGCCs in generating a wide kinetic range of Ca^2+^ spikes in CA3PCs.

**Figure 5.**
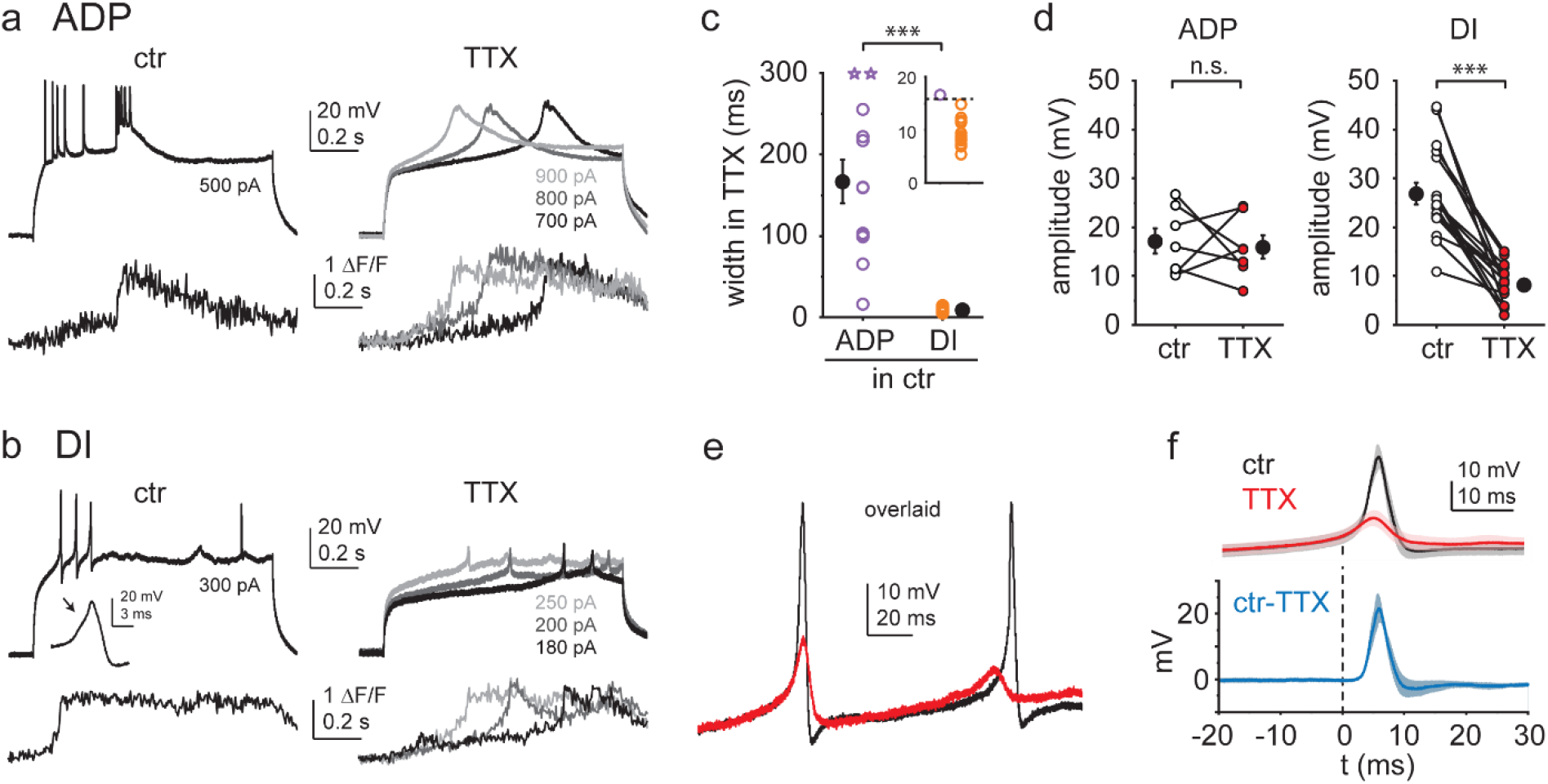
Ca^2+^ spikes with different kinetics are mediated by different ionic mechanisms. **a-b)** Representative experiments showing the effect of 1 μM TTX on ADPs (a) and DI spikes (b). V_m_ (top) and distal dendrite Ca^2+^ signals (bottom) under control conditions (left) and after application of TTX at various I_inj_ levels. **c)** Width of TTX-resistant Ca^2+^ spikes in cells with ADP only (purple, n=12) and cells expressing DI spikes (orange, n=13). Open circles, individual dendrites (open stars: dendrites where width was maximized as 300 ms because V_m_ did not return to half amplitude within the duration of the I_inj_ step); filled black symbols, mean ± SEM. Inset shows the 0-20 ms width range magnified. **d)** Comparison of spike amplitude before and after TTX application in dendrites with ADPs only (left, n=7) and dendrites with DI spikes (right, n=17). **e)** Overlaid voltage traces from a dendrite in control (black) and after TTX application (red). **f)** Summary of the impact of TTX on DI spike kinetics Top, V_m_ traces before (black, ctr) and after (red) TTX application (mean ± SEM of 6 experiments), aligned to 1 V/s. Bottom, result of the subtraction of ctr and TTX traces. The fast TTX-sensitive component follows the initial slow depolarization. **Fig. 5 – figure supplement 1. Additional properties of dendritic Ca^2+^ spikes**

To our surprise, we noticed that the amplitude (and dV/dt) of TTX-resistant fast spikes was consistently smaller than that of the DI spikes in ACSF (in ACSF: 26.9 ± 2.2 mV, in TTX: 8.2 ± 1.0 mV, n=17, Wilcoxon test, p<0.001), whereas the amplitudes of ADPs and TTX-resistant slow spikes were not systematically different (17.2 ± 2.6 mV before and 15.9 ± 2.4 mV after TTX application, n=7, p=0.735, Wilcoxon test, **Fig. 5d**). Therefore, we examined in more detail the possibility that VGNCs contribute in some form selectively to DI spikes. First, we determined which parameters of the spike were most affected by TTX. Our analysis showed that the reduction of spike amplitude was largely caused by a drop in peak voltage (from −0.8 ± 2.2 mV to −15.6 ± 1.3 mV, n=17, p<0.001, Wilcoxon test, **Fig. 5 – figure supplement 1d**) rather than the modest increase in spike threshold (from −27.2 ± 1.0 mV to −23.7 ± 1.0 mV, n=17, p<0.01, Wilcoxon test, **Fig. 5 – figure supplement 1e-f**). Furthermore, overlaying individual traces (**Fig. 5e**) or aligning averaged voltage responses in control and TTX to a specific dV/dt value (1 V/s; **Fig. 5f**) revealed that TTX did not affect the initial slow, concave rise of the spike, but exclusively reduced the subsequent, fast-rising peak component. As an independent confirmation of the involvement of VGNCs in the generation of DI spikes, replacement of a majority of extracellular Na^+^ ions in the ACSF with the large cation NMDG^+^ (which permeates less through VGNCs) produced a similar effect to that of TTX (**Fig. 5 – figure supplement 1c, g**). Altogether, these results suggest that DI spikes can be mediated by a unique hybrid mechanism, whereby initiation of a fast Ca^2+^ spike can recruit regenerative activation of VGNCs that further amplify and sharpen the voltage response, with the two components blending smoothly into a rapid and transient combined dendritic spike.

### Different Ca^2+^ spike types obey distinct compartmentalization rules

The distinct characteristics of fast and slow Ca^2+^ spike components raise the question whether their propagation and compartmentalization properties are also different. To address this, we recorded Ca^2+^ spike-associated dendritic Ca^2+^ signals (in TTX) both distally within the same dendritic family (250 ± 28 μm distal from the patch pipette, n=14 cells; depicted as d1 in **Fig. 6a**) and in another dendrite branching off more proximally, typically from a different low-order trunk segment (d2; dendritic distance from pipette: 294 ± 23 μm, n=14, **Fig. 6a**). We first used the high-affinity dye OGB-1 to be able to detect even small increases in Ca^2+^ as a reporter of spike propagation within and across dendritic compartments. We found a strong difference between fast and slow spikes in their propagation properties (2-way repeated measures ANOVA, p<0.001 for kinetic group, p=0.005 for location, p<0.001 for interaction). Slow Ca^2+^ spikes (width > 60ms) were accompanied by large (likely dye-saturating) Ca^2+^ signals both in d1 and d2 (**Fig. 6b-c**, n=5 cells, p=0.781, Tukey’s post hoc test), and even in basal dendrites (measured in n=3 cells, **Fig. 6d**), indicating efficient propagation across major bifurcation points or even the entire dendritic tree. In contrast, fast Ca^2+^ spikes (width <20 ms) were restricted to the dendritic family connected to the patched trunk (**Fig. 6b-c**; n=9 cells, p<0.001, Tukey’s post hoc test). Finally, to assess the local propagation capacity of fast Ca^2+^ spikes in more detail, we mapped fast Ca^2+^ spike-associated Ca^2+^ signals in higher spatial resolution proximally and distally from the patch pipette (using the low-affinity Ca^2+^ dye OGB-6F to reduce complications from dye saturation). These experiments revealed relatively uniform, large Ca^2+^ signals towards distal dendritic locations, but a strong drop of Ca^2+^ signals within ∼160 μm from the pipette in the proximal direction, so that fast spikes evoked in a higher order trunk did not propagate closer than ∼100 μm to the soma (**Fig. 6d-f**, n=4 experiments).

**Figure 6.**
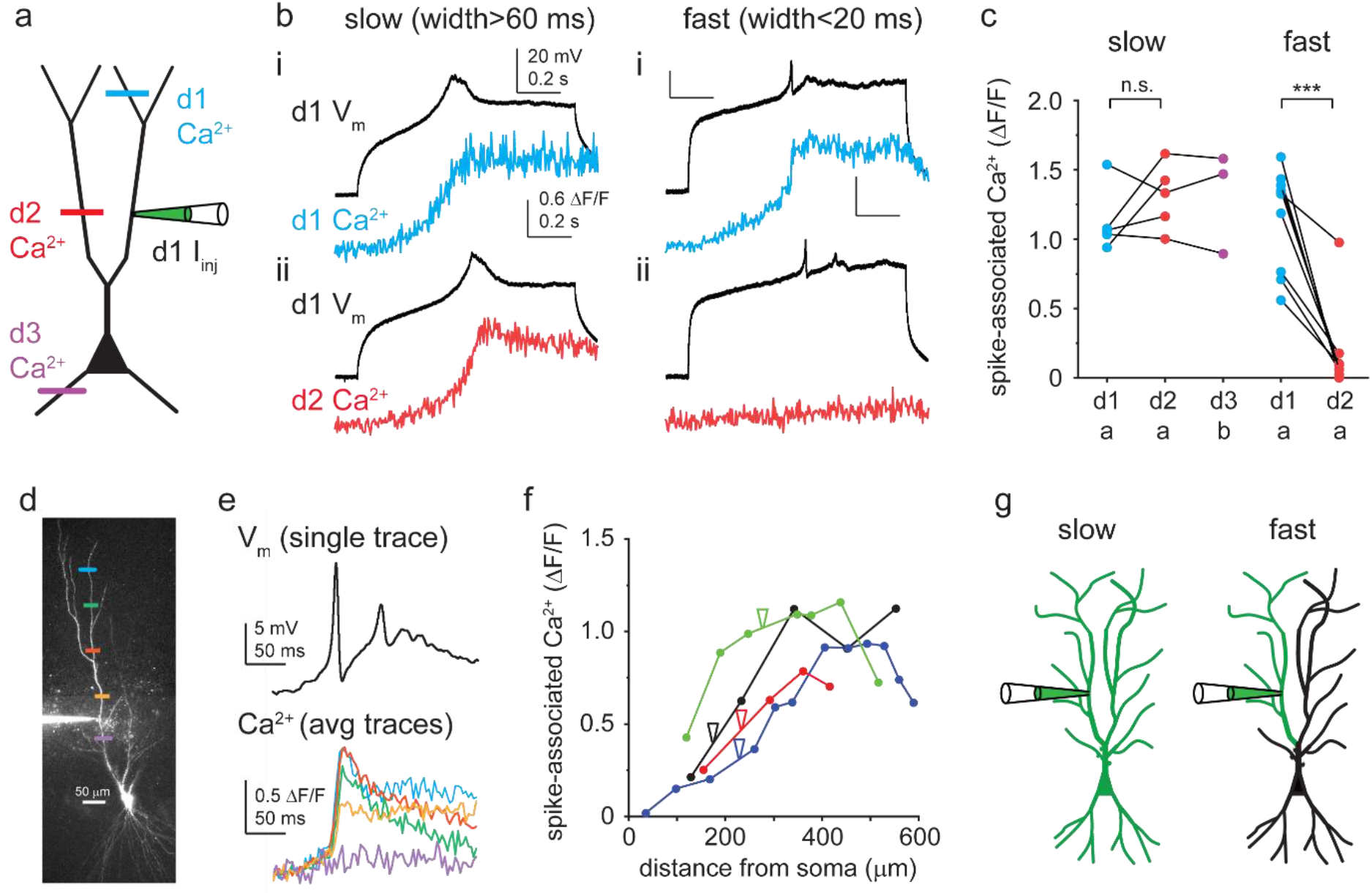
Propagation of different Ca^2+^ spike types. **a)** Schematic of experimental strategy. Dendritic Ca^2+^ signals were measured as a proxy to assess Ca^2+^ spike propagation in different regions of the arbor. **b)** Representative TTX-resistant slow (left) and fast (right) Ca^2+^ spikes. Spike-evoked Ca^2+^ signals (OGB-1) measured in the patched (i) and in a different (ii) apical dendritic subtree. **c)** Summary of spike-evoked Ca^2+^ signal amplitudes at different dendritic tree parts. Note that large spike-evoked Ca^2+^ signals likely saturate OGB-1. **d)** 2P stack of a CA3PC, with Ca^2+^ measurement sites indicated by colored lines. **e)** Fast Ca^2+^ spike-evoked Ca^2+^ signals (measured with OGB-6F) at the locations indicated in panel d. Ca^2+^ signals were aligned to spike onset (one trace shown on top). **f)** Distance-dependence of fast Ca^2+^ spike-associated Ca^2+^ signals in 4 experiments. Note the drop of Ca^2+^ signals from the pipette towards the soma. **g)** Concept of compartmentalization rules. Slow Ca^2+^ spikes are global events, whereas fast Ca^2+^ spikes are restricted to apical dendritic subtrees.

These results indicate different generation and propagation properties of different Ca^2+^ spike types (**Fig. 6g**). Slow Ca^2+^ spikes are apparently global events that engage virtually the entire dendritic tree. In contrast, fast Ca^2+^ spikes are generated within the dendritic family of the stimulated trunk, and although they propagate well distally, they cannot efficiently invade the proximal main or sibling trunk segments, creating a mesoscale compartmentalization level represented by separate apical dendritic subtrees.

### Morphological correlates of distinct active dendritic properties

The dendritic structure of CA3PCs is highly diverse, and their electrophysiological properties were proposed to be related to their topographic position within the CA3 area and dendritic morphology (Bilkey and Schwartzkroin, 1990; Ding et al., 2020; Hunt et al., 2018; Raus Balind et al., 2019; Sun et al., 2017). A recent study even suggested the existence of a sparse class of CA3PCs with bursting phenotype that lack thorny excrescences (TEs) and input from mossy fibres (“athorny” cells, Hunt et al., 2018). Therefore, we examined the relationship between Ca^2+^ spike phenotype and morphological properties of CA3PCs in our dataset. In almost all of our fluorescently labelled cells we unambiguously confirmed the presence of TEs. The estimated coverage of the proximal trunk(s) by TEs varied widely (**Fig. 7a-b**) and was inversely correlated with the length of first-order apical trunk(s) (**Fig. 7c**, Spearman correlation R=-0.495, p<0.001, n=89), confirming previous studies (Fitch et al., 1989) and suggesting a gradient of morpho-functional properties of regular CA3PCs. We found that the dendritic Ca^2+^ spike profile was correlated with these basic anatomical features of CA3PCs. First, CA3PCs expressing DI spikes in the recorded dendrite (with or without ADPs) had on average twice as high TE coverage (**Fig. 7d**, ADP-only: 40 ± 5 μm, n=19; DI: 84 ± 8 μm, n=35, Mann-Whitney test: p<0.001) than ADP-only cells. Second, cells with DI spikes had shorter primary trunks (**Fig. 7e**, ADP-only: 128 ± 11 μm, n=20; DI: 57 ± 5 μm, n=41, Mann-Whitney test: p=0.002) and more often had multiple first-order trunks (**Fig. 7f**, ADP-only: 1 ± 0, n=20; DI: 1.49 ± 0.11, n=41, Mann-Whitney test: p<0.001) than ADP-only cells. Interestingly, although the distance of the dendritic patch pipette from the soma was similar in the two electrophysiological groups (ADP-only: 276 ± 15 μm, n=20; DI: 260 ± 10 μm, n=41, Mann-Whitney test: p=0.282, see also **Fig. 3h**), the dendritic path between the pipette and soma contained more branch points in the case of dendrites with DI spikes (**Fig. 7g**, ADP-only: 4.15 ± 0.36, n=20; DI: 5.97 ± 0.35, n=39, Mann-Whitney test: p<0.001). This may be consistent with the idea that dendrites with more branch points are electrically more isolated from the soma (Vetter et al., 2001). In summary, the morphological diversity of CA3PCs corresponds to distinct dendritic excitability phenotypes.

**Figure 7.**
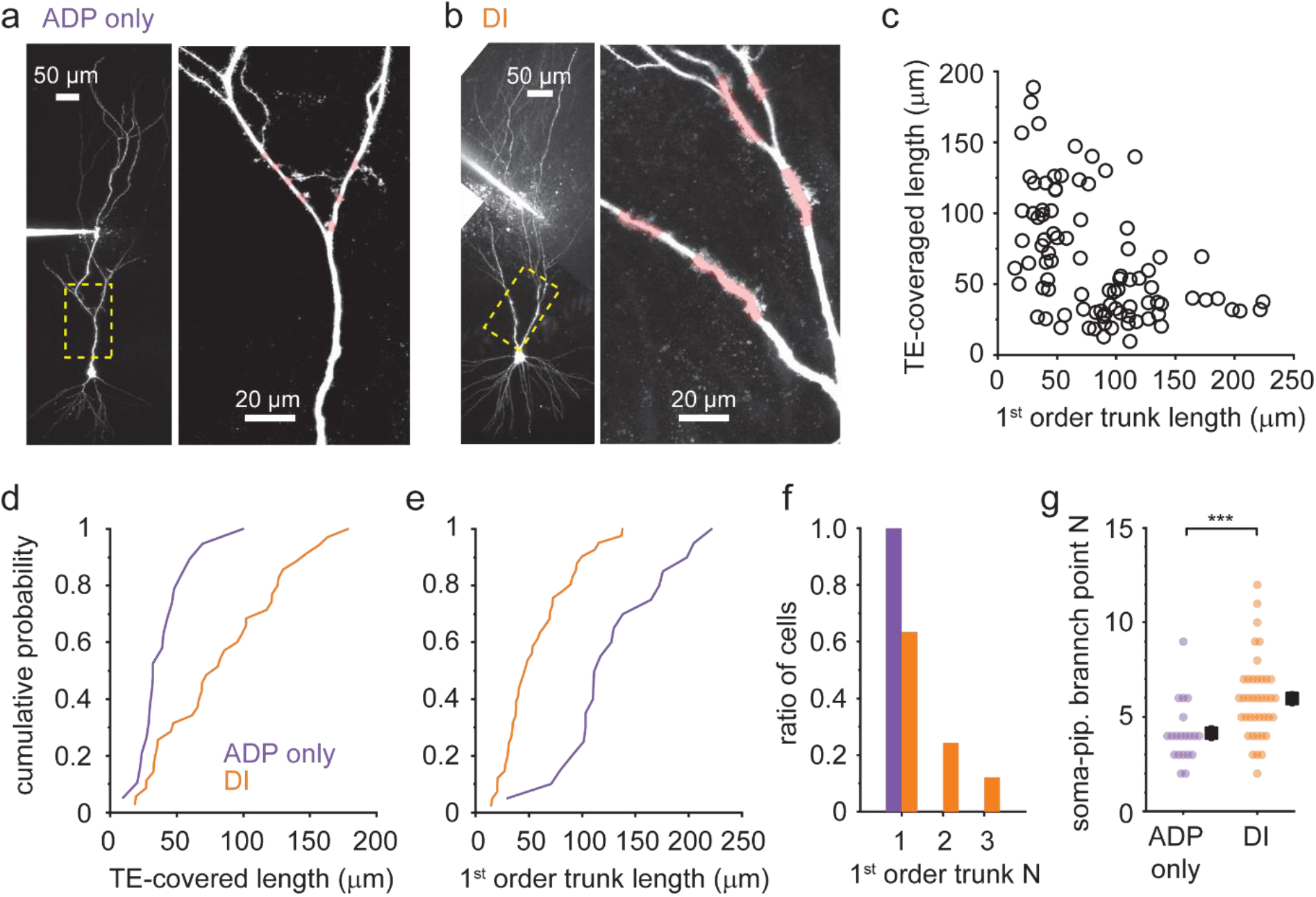
Dendritic Ca^2+^ spike phenotype correlates with morphological traits. **a-b**) Representative 2P stack of two CA3PCs with ADP-only (**a**) or DI (**b**) Ca^2+^ spike phenotypes (with ≤600 or 700pA I_inj_). Yellow dashed boxes are enlarged on the right panels. Trunk segments with TEs are indicated by pink shading. **c**) Relationship between first-order apical trunk length and total TE-covered dendrite length among CA3PCs (n=89). For cells with multiple primary trunks the mean trunk length is shown. **d-e**) Cumulative probabilities of total TE-covered dendrite length (**d**, n=19 ADP-only, n=35 DI) and first-order apical trunk length (**e**, n=20 ADP-only, n=41 DI) for cells with dendritically recorded ADP-only and DI Ca^2+^ spike types. **f**) Number of first-order apical trunks for the two electrophysiological groups (n=20 ADP-only, n=41 DI). **g**) Number of branch points between the patch pipette and the soma in the two electrophysiological groups.

### Distinct Ca^2+^ spike types have opposing impact on the form of somatic output

How do the diverse dendritic Ca^2+^ spike types influence the somatic output of CA3PCs? The widely accepted idea is that the primary effect of Ca^2+^ spikes on output is the generation of bursts. ADP-type Ca^2+^ spikes indeed served such a role: they often triggered additional somatic spike(s) to produce short (2-3 APs) or long (>3 APs) series of APs or bursts after the first initiating AP (**Fig. 8a, b**). In contrast, DI spikes never evoked more than a single somatic AP (**Fig. 8a-c**). Furthermore, compared to simple bAPs (s-APs), the bAP evoked by DI spikes (Ca-APs) was followed by a faster repolarization rate (ratio of dV/dt_min_ (Ca-APs/s-APs): 1.29 ± 0.04, p=0.001, Wilcoxon-test compared to 1) and enhanced fast afterhyperpolarization (difference of AHP V_min_ (Ca-AP - s-AP): −2.68 ± 0.34 mV, p=0.001, Wilcoxon test compared to 0) in the dendrites (n=13, pooled dual (n=5) and dendrite-only (n=8) recordings, **Fig. 8c-e,** and **Fig. 8 – figure supplement 1**) as well as larger AHP at the soma (s-AP: −3.21 ± 0.89 mV, Ca-AP: −4.70 ± 1.12 mV, measured in n=5 dual recordings, p=0.043, Wilcoxon-test, **Fig. 8c, d, f**). Thus, the impact of DI spikes on somatic output is the opposite of that classically proposed for dendritic Ca^2+^ spikes: they not only trigger strictly single APs, but they even *suppress* the probability of consecutive AP generation and thereby actively reduce burst output.

**Figure 8.**
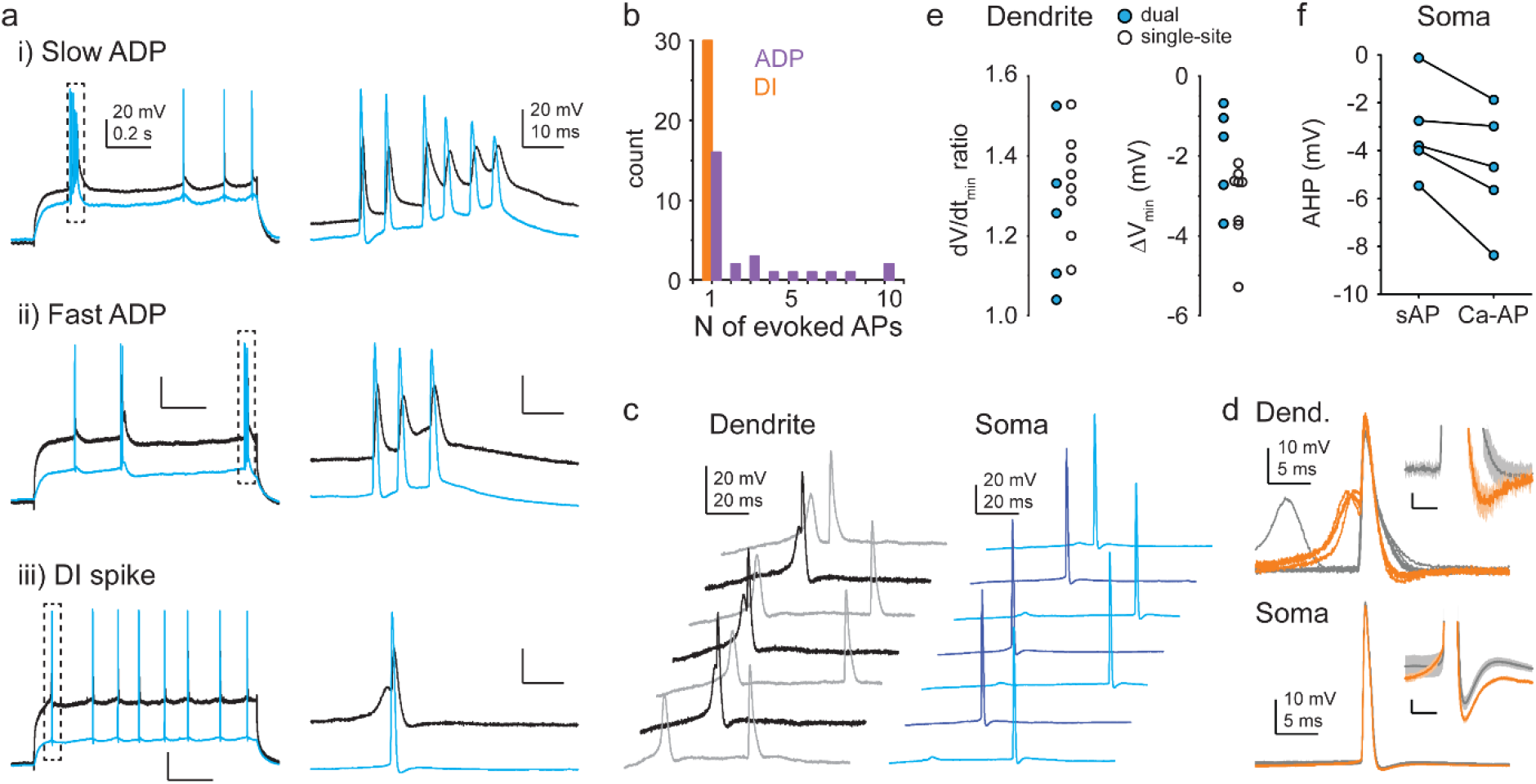
Opposing impact of different Ca^2+^ spike types on somatic output. **a)** Example of AP output evoked by slow ADP (top), fast ADP (middle) and DI spike (bottom) in dual recordings (3 different cells). **b)** Median number of APs evoked at threshold I_inj_ by ADPs and DI spikes in individual cells. Note that n=1 AP evoked by an ADP results in a burst of 2 APs together with the AP initiating the ADP. Only cells with at least three Ca^2+^ spike events evoking APs were included in the analysis. **c)** Example traces from a dual recording. Darker colors indicate traces with Ca-APs. **d)** Ca-APs (orange) and s-APs (grey) aligned to peak from the experiment shown in c. Insets show AHPs enlarged; scale bars: 1 mV, 3 ms. **e)** Ratio of repolarization rate (dV/dt_min_, left) and difference in minimum V_m_ within 12 ms after peak (right) of Ca-APs and s-APs measured in dendrites. Blue-filled circles: dual recordings; open circles: dendrite-only recordings. f) Somatic AHP amplitude following Ca-APs and s-APs in dual recordings. **Fig. 8 – figure supplement 1. Effect of DI spikes on AP output in single-site dendritic recordings**

## Discussion

Using dual soma-dendritic and single-site dendritic patch-clamp recordings combined with 2P Ca^2+^ imaging, we reveal complex active properties of CA3PC apical dendrites. Besides supporting local Na^+^ and NMDAR-mediated d-spikes (Brandalise and Gerber, 2014; Kim et al., 2012; Makara and Magee, 2013), we report that these dendrites express distinct types of Ca^2+^ spikes that can oppositely impact the form of firing by the neuron.

One prominent component is provided by slow Ca^2+^ spikes, which are primarily responsible for generating ADP following initial AP(s) and evoke bursts of additional APs at the soma. While ADP-type Ca^2+^ spikes have been described in apical trunks of various PC types, their properties have rarely been systematically evaluated and compared. We found that the slow Ca^2+^ spikes underlying ADPs in CA3PCs are typically longer lasting (on average ∼3-fold) than similar type Ca^2+^ spikes in CA1PC trunks. We also showed that slow Ca^2+^ spikes appear globally throughout the apical (or even the whole) dendritic arbor, and they are most prevalent in CA3PCs with relatively long single primary trunks. These properties altogether well align with the high burst propensity observed in deep distal CA3PCs, which show this morphological feature (Hunt et al., 2018; Raus Balind et al., 2019). At this point it is not clear whether slow Ca^2+^ spikes are evoked *ab ovo* as a “global” dendritic spike (Connelly et al., 2015), or they have a specific generation zone from where they invade the arbor. The results suggest that the long primary apical trunk acts as a designated generation site for slow Ca^2+^ spikes, but it is also possible that this dendritic trait simply correlates with other features (possibly including passive and/or active ion channel-mediated dendritic properties) that promote slow spike generation.

In addition to slow Ca^2+^ spikes, depolarization also evoked a novel fast form of dendritically initiated Ca^2+^ spikes in a large fraction of CA3PC trunk dendrites. DI spikes are generated by fast Ca^2+^ spikes, which, if large enough, can recruit subsequent contribution by regenerative VGNC activation. Especially when combined into such hybrid d-spike, fast Ca^2+^ spikes can produce large enough depolarization in a window above threshold stimulation to efficiently evoke a tightly coupled AP at the soma. Curiously, these APs are immediately followed by enhanced repolarization and AHP both at dendrite and soma, cutting off depolarization and actively blocking burst firing. As a result, DI spikes produced strictly single APs, a form of input-output transformation that is the opposite of that generally proposed for Ca^2+^ spikes. This means that the regular spiking firing profile of certain CA3PCs may be not simply due to a lack of d-spikes (or other mechanisms) promoting bursts, but instead due to a specific form of local dendritic spikes that actively produces non-bursting, regular spiking firing phenotype. From another point of view, DI spikes might serve to amplify certain local apical synaptic input forms without strong perisomatic dendritic activity (i.e. no bAPs) to promote an output, yet they do so in a way that prevents switching to burst firing, which can be preserved for signaling different input patterns, e.g. associative input conjunction (Raus Balind et al., 2019).

Another interesting aspect of DI spikes is their semi-compartmentalization: they propagate well towards distally in the dendritic subtree belonging to the depolarized intermediary trunk, but they propagate poorly proximally and fail near the major bifurcation zone ∼100 μm from the soma. Thus, dendritic families composed of higher-order trunks with connected daughter branches can correspond to independent spatial units of synaptic integration and/or plasticity (∼3-5 units per cell). This represents a ‘mesoscale’ level of compartmentalization that is intermediate between global and local (branch specific) d-spikes. One might speculate that DI spikes could be particularly suited to promote compartmentalized induction of synaptic or intrinsic (Losonczy et al., 2008) plasticity, because the evoked single APs, backpropagating from the soma to other parts of the dendritic arbor, would likely be inefficient to induce substantial plasticity outside the spike-generating dendritic family. Such compartmentalized plasticity could lead to biased wiring of correlated synaptic inputs targeting specific apical subtrees. It remains to be seen whether such mesoscale connectivity structure exists in CA3PCs, in addition to the fine-scale clustering of inputs previously shown (Takahashi et al., 2012). Notably, due to their strong attenuation, DI spikes are challenging to detect and likely remain overlooked with somatic voltage recording techniques; multisite measurements in dendrites and soma, e.g. with fast voltage imaging techniques will be required to uncover this form of input-output transformation *in vivo*.

While we classified the dendrites based on whether they expressed DI spikes, it is likely that large transient ADPs that we also observed in some of the dendrites were mediated by a similar fast Ca^2+^ spike mechanism to that producing DI spikes. In fact, in several dendrites smaller I_inj_ evoked fast ADPs whereas at higher I_inj_ DI spikes occurred. We speculate that at weaker depolarization the arrival of a bAP can provide the necessary stimulus to trigger the Ca^2+^ spike that therefore appears as an ADP; the inactivation of VGNCs during the initializing bAP prevents their further contribution to the spike. In contrast, stronger depolarization can itself initiate the fast Ca^2+^ spikes, which in turn can trigger an additional Na^+^ spike component to develop the full fast hybrid DI spike. Notably, a substantial fraction of CA3PCs co-expressed different types of Ca^2+^ spikes. Altogether, the various forms of fast and slow dendritic Ca^2+^ spikes in the complex branching apical tree provide particularly large room for specific input-output transformations in CA3PCs depending on the strength and actual spatiotemporal pattern of activity of the three different sources of afferent synaptic inputs.

Further studies are required to elucidate the biophysical mechanisms and implications of the diverse spike characteristics. In particular, future work is needed to resolve whether the diversity may be explained by differences in the molecular composition or functional properties of various VGCC subtypes producing kinetically different Ca^2+^ currents (Avery and Johnston, 1996) or other, particularly K^+^ channel types (e.g. specific voltage-dependent or Ca^2+^-activated K^+^ channels), as well as the role of passive properties. While ion channels may be regulated in an activity- or state-dependent fashion, the observed correlation of functional and morphological properties of CA3PCs suggests an at least partially rigid organization of d-spike mechanisms in different dendritic compartments (although proximal apical CA3 dendritic morphology is plastic (Juraska et al., 1989; McEwen and Magarinos, 1997)). A reasonably long primary trunk appears to allow the generation of slow Ca^2+^ spikes with a low threshold that can be reached both by synaptic/dendritic and somatic depolarization (Raus Balind et al., 2019), whereas higher-order trunks may be diverse individual Ca^2+^ spike generators. It needs to be studied by future work how the summation of the various d-spikes at the soma finally determines the shape and burstiness of AP output. It will be also interesting to explore the theoretical aspects of how the complex rules driving specific output responses and plasticity mechanisms evoked by different state-dependent input combinations can contribute to the postulated pattern separation and pattern completion functions of the CA3 network during spatial navigation and memory processes.

The functional properties of DI spikes we discovered in rat CA3PCs resemble those recently described for dendritic Ca^2+^ spikes (called dCaAPs) in L2/3PCs of the human neocortex derived from patients operated for epilepsy or tumour (Gidon et al., 2020). Except for somewhat different amplitudes, the kinetics and the inverse relationship of amplitude with depolarization of these spikes in rat CA3PCs was similar to that in hL2/3PCs. While dCaAPs of human neurons were present in TTX (similar to our results), comparison of spike properties before and after TTX application was not reported; therefore, it remains to be determined whether dCaAPs in human neurons have similar hybrid Ca^2+^/Na^+^ spike components as DI spikes in rat CA3PCs. Altogether, our results contradict the idea that DI Ca^2+^ spikes would have developed to support human specific cortical computations. Instead, it is intriguing to speculate that DI spikes may contribute to basic circuit computation motifs shared by hippocampal CA3 and superficial cortical PCs, for example related to their intermediate position in a sequential chain of input processing (Shepherd, 2011). Finally, the presence of DI Ca^2+^ spikes in neurons of healthy rats is a strong indication for their physiological role and argues against a possibility that the expression of these spikes in human neurons could have been related to disease or pharmacological treatment.

Altogether, our results point out that different types of PCs may utilize different structure-function models for dendritic computations (i.e. different architecture of hierarchical and parallel nodes of nonlinear synaptic integration) that allow complex forms of input-output transformations. The cell type specific forms of dendritic spikes raise the idea that active dendrites may support circuit-specific computations, whereas the heterogeneity within a principal cell class suggests that PC subpopulations may be dedicated to perform different information processing functions. Elucidating how these diverse models are employed *in vivo* under behaviorally relevant conditions of natural complex excitatory input patterns, inhibition and neuromodulation (Sheffield et al., 2017; Takahashi et al., 2020) will be a major step towards understanding how single cell computations can serve the network functions underlying appropriate and flexible behaviors to adapt to environmental challenges.

## Materials and Methods

### Hippocampal slice preparation

Adult male Wistar rats (7-12-week-old) were used to prepare 400-µm-thick slices from the hippocampus as described (Makara and Magee, 2013; Raus Balind et al., 2019), according to methods approved by the Animal Care and Use Committee of the Institute of Experimental Medicine, and in accordance with the Institutional Ethical Codex, Hungarian Act of Animal Care and Experimentation 40/2013 (II.14), and European Union guidelines (86/609/EEC/2 and 2010/63/EU Directives). Animals were deeply anaesthetized with 5% isoflurane and quickly perfused through the heart with ice-cold cutting solution containing (in mM): sucrose 220, NaHCO_3_ 28, KCl 2.5, NaH_2_PO_4_ 1.25, CaCl_2_ 0.5, MgCl_2_ 7, glucose 7, Na-pyruvate 3, and ascorbic acid 1, saturated with 95 % O_2_ and 5 % CO_2_. The brain was quickly removed and slices were prepared in cutting solution using a vibratome (VT1000A, Leica, Leica Biosystems GmbH, Nussloch, Germany). Slices were incubated in a submerged holding chamber in ACSF at 35 °C for 30 min and then stored in the same chamber at room temperature.

### Patch-clamp recordings

Slices were transferred to a custom-made submerged recording chamber under the microscope where experiments were performed at 32-34 °C in ACSF containing (in mM): NaCl 125, KCl 3, NaHCO_3_ 25, NaH_2_PO_4_ 1.25, CaCl_2_ 1.3, MgCl_2_ 1, glucose 25, Na-pyruvate 3, and ascorbic acid 1, saturated with 95 % O_2_ and 5 % CO_2_. Neurons were visualized using Zeiss Axio Examiner or Olympus BX-61 epifluorescent microscope under infrared illumination and water immersion lens (63X or 60X during recording, 20X or 10X for overview z-stacks, Zeiss or Olympus).

Higher order apical dendritic trunks in str. radiatum were patched under oblique illumination at 268 ± 7 μm distance from the soma. After establishing the dendritic whole-cell configuration, cells were loaded for >10 minutes to visualize the soma and dendritic tree. Neurons were carefully inspected for thorny excrescences on thick parent dendrites (to verify they were CA3PCs), and that no main dendritic trunk was cut. In dual recordings, the soma was patched subsequently, either under oblique illumination or guided by 2P imaging. Neurons included in this study were all recorded in different slices, typically from different animals.

Dendritic (6-10 MΩ) and somatic (4-6 MΩ) patch pipettes were filled with a solution containing (in mM): K-gluconate 134, KCl 6, HEPES 10, NaCl 4, Mg_2_ATP 4, Tris_2_GTP 0.3, phosphocreatine 14 (pH=7.25) complemented with 50 µM Alexa Fluor 594 and 100 µM Oregon Green BAPTA-1 (OGB-1) or Oregon Green BAPTA-6F (OGB-6F) (all fluorescent dyes from Invitrogen-Molecular Probes). Electrophysiological results were similar using OGB-1 and OGB-6F and therefore results obtained with different Ca^2+^-sensitive dyes were pooled.

Current-clamp whole-cell recordings were performed using BVC-700A amplifiers (Dagan, Minneapolis, MN, USA) in the active ‘bridge’ mode, filtered at 3 kHz and digitized at 50 kHz. Series resistance was typically between 15-30 MΩ at the soma and 25-60 MΩ at the dendrite (if possible, reduced by gently blowing into the pipette), frequently checked and compensated with bridge balance and capacitance compensation. Membrane potentials (V_m_) are reported without correction for liquid junction potential (∼10 mV). The baseline V_m_ was kept at -68- - 72 mV with appropriate constant current injection. Dendritic V_m_ of CA3PCs ranged between - 53 to -73 mV, R_in_ was 105 ± 3 MΩ. In all dual recordings, somatic V_m_ was more negative than −60 mV (Raus Balind et al., 2019). A set of experiments were performed on apical trunk dendrites of CA1PCs at ∼230-460 μm dendritic distance from the soma; experiments were identical to those in CA3PCs except that V_m_ was held between -64 - -68 mV. Dendritic V_m_ of CA1PCs ranged between −59 to −62 mV, and R_in_ was 44 ± 9 MΩ.

TTX, NBQX disodium salt, D-AP5 (Tocris) and NiCl_2_ (Sigma) were prepared in stock solution in distilled water, stored at −20 °C (NiCl_2_ at room temperature) and dissolved to final concentration into bubbled ACSF before application. To reduce extracellular Na^+^ concentration, in some experiments NaCl was replaced with equal concentration of NMDGCl in the ACSF. Modified ACSF solutions were applied for at least 10 minutes before testing their effect.

### Two photon imaging and uncaging

A dual galvanometer based two photon scanning system (Bruker, former Prairie Technologies, Middleton, WI, USA) was used to image the patched neurons and to uncage glutamate at individual dendritic spines as previously described (Makara and Magee, 2013; Raus Balind et al., 2019). Two ultrafast pulsed laser beams (Chameleon Ultra II; Coherent, Auburn, CA, USA) were used: one laser at 920 or 860 nm for imaging OGB dyes and Alexa Fluor 594, respectively, and the other laser tuned to 720 nm to photolyze MNI-caged-L-glutamate (Tocris; 10 mM in ACSF) that was applied through a puffer pipette with a ∼20-30- μm-diameter, downward-tilted aperture above the slice using a pneumatic ejection system (PDES-02TX (NPI, Tamm, Germany). The intensity of the laser beams was independently controlled with electro-optical modulators (Model 350-80, Conoptics, Danbury, CT, USA). Linescan Ca^2+^ measurements were performed with 6-8.8 μs dwell time at ∼200-300 Hz.

Glutamate uncaging was performed at a clustered set of 20 spines on the patched apical trunk, using 0.5 ms uncaging duration at each spine with 0.1 ms intervals between synapses, repeated 5 times at 40 Hz (i.e. gamma burst stimulus)(Raus Balind et al., 2019). Uncaging laser power was adjusted to yield compound voltage responses near the threshold of regenerative events (bAPs or dendritic spikes), preferably so that both subthreshold and suprathreshold responses could be evoked. The average amplitude of the voltage response at the first pulse was 19.2 ± 2 mV (n=10 dendrites).

### Data analysis

Analysis of voltage and Ca^2+^ recordings was performed using custom-written macros in IgorPro (WaveMetrics, Lake Oswego, OR, USA). For calculating input resistance and soma-dendrite voltage transfer, we measured the steady-state voltage change to 50 pA, 300 ms hyperpolarizing step current injections from baseline V_m_. In cells with a high rate of spontaneous EPSPs the above electrophysiological properties were not determined.

Ca^2+^ signals are expressed as ΔF/F_0_ = (F(t)-F_0_)/F_0_, where F(t) is fluorescence at a given time point and F_0_ is the mean fluorescence during 50 ms preceding the depolarizing I_inj_. To measure Ca^2+^ spike associated Ca^2+^ signal amplitude (Fig. 6), traces were aligned to the initial rise (300-400 mV/s dV/dt value) of the Ca^2+^ spike, and we calculated the difference between the maximum average of 10 (OGB-1) or 3 (OGB-6F) consecutive points following the spike and the average of the 20-50 ms period preceding the spike. Note that this measurement may slightly overestimate the amplitude. Ca^2+^ traces and some of the electrophysiological traces were slightly smoothed (binomial filter, N=1) for display purposes only.

Ca^2+^ spike properties were quantified using I_inj_ levels at or 100pA above the lowest I_inj_ evoking the spike, unless otherwise indicated. Usually 5-10 traces were recorded with each I_inj_ level, and data were averaged. Where multiple Ca^2+^ spikes occurred within a trace, we analyzed the one with the largest amplitude, which was most often the first event.

ADPs were defined as regenerative depolarization initiated by 1 or 2 preceding APs. Fast ADP amplitude was measured on events which showed additional rising phase and clear peak following the initiating AP(s). We avoided ADP amplitude measurement in cases where the peak could not be defined (such as with constant decay of voltage or due to the generation of a consecutive AP) and when the ADP-evoking AP was preceded by >2APs in the burst. ADP peak amplitude was measured from the V_m_ immediately preceding the first AP initiating the ADP. We note that this measurement likely overestimates the true Ca^2+^ spike amplitude due to the tail of the decaying AP on which the Ca^2+^ spike is riding. Slow ADPs did not have a clear peak; amplitude was measured as the maximum sustained depolarization between consecutive APs in the burst compared to the V_m_ immediately preceding the first initiating AP. Width at half amplitude of ADPs was not determined due to the uncertain rise and amplitude.

Dendritically initiated (DI) spikes were defined as regenerative events that were not initiated by APs. Peak amplitude of the DI spike was determined in cases where voltage reached a clear peak (either alone or before evoking a consecutive AP). The threshold of DI spikes was measured at the inflection point where voltage began to deviate from subthreshold baseline depolarization. This was sometimes difficult to determine, and when possible (at I_inj_ experiments) it was aided by extrapolating to the baseline where voltage returned to after the spike, or by fitting a double exponential to the initial subthreshold voltage response to the step current injection. Width (at half amplitude) of DI spikes was measured on smoothed traces (binomial filter, N=10) on spikes with at least 5 mV amplitude and when no APs were evoked. The parameters of pharmacologically isolated Ca^2+^ spikes (measured in TTX or NMDG), were determined the same way as those of DI spikes.

The dV/dt ratio, used to support distinguishing Ca^2+^ spikes and simple APs (Fig. 1 – figure supplement 1), was defined for each dendritic regenerative event as dV/dt_pre_/dV/dt_peak_, where dV/dt_peak_ is the maximum dV/dt value during the event, and dV/dt_pre_ is the maximum dV/dt value in the 1.5-9 ms time window preceding dV/dt_peak_.

To quantify the number of evoked APs by suprathreshold Ca^2+^ spikes (Fig. 8b), results obtained at the threshold level of I_inj_ (and if not enough events, at threshold +100pA) evoking Ca^2+^ spikes were analyzed, in those experiments where at least 3 Ca^2+^ spike events (ADP or DI spike) with evoked APs were recorded. DI spikes were included in the analysis only if they were clearly separable from the evoked AP. ADP events were analyzed only after steady-state voltage was reached by I_inj_. APs following ADPs were considered to be evoked by the Ca^2+^ spike if they occurred before the depolarized V_m_ returned to 3.5 mV above the baseline but maximum within 300 ms time window after the first AP, with <100 ms preceding interspike interval.

When measuring the impact of DI spikes on the AHP (Fig. 8), in order to ensure that temporal voltage changes did not affect our comparison, we restricted our analysis to those cells where both Ca^2+^ spike-evoked APs (AP_Ca_) and simple APs (AP_s_) were evoked at the same I_inj_ level, either within the same traces at the steady-state depolarization phase, or on intermittent traces within the same time window.

To calculate the propensity of dendritic spikes evoked by 2PGU, traces with moderate stimulus strength were used, i.e. where regenerative events (bAPs and/or d-spikes) were evoked only on any of the last 3 pulses of the gamma burst. The relative ADP and DI spike probability (range: 0-1) was calculated by dividing the number of traces displaying the respective d-spike type by the total number of suprathreshold traces (minimum n=3).

Transfer of steady-state voltage signals in dual recordings was determined using hyperpolarizing current injection either to the soma or dendrite, and calculating the ratio of the resulting voltage deflection at the two locations. Attenuation of dendritic spikes was measured as the ratio of amplitudes at the dendrite and the soma. The somatic peak of fast ADPs was in some cases masked by the AHP of the AP; these cases were not included in the analysis.

We note that the number of measurements varies for different parameters due to the specific criteria applied for their analysis.

### Morphological analysis

Alexa Fluor 594 fluorescence was used for morphological analysis. Dendritic morphological and distance measurements were performed using ImageJ (NIH, Bethesda, MD, USA) on stacked images collected at the end of the experiment. Dendritic length and distance were measured on the collapsed 2P stacks by manually drawing a segmented line along the dendrite. Thorny excrescences were identified as large, lobular, complex spine-like postsynaptic structures on first- or higher order trunks near the soma (Chicurel and Harris, 1992; Raus Balind et al., 2019). The total coverage of dendrites by thorny excrescences was estimated as the sum of the length of freehand lines drawn manually along the dendritic segments where thorny excrescences were observed.

### Statistical analysis

No statistical methods were used to predetermine sample sizes, but our samples are similar to or exceed those reported in previous publications and that are generally employed in the field. Statistical analysis was performed with the Statistica software (Statsoft, Tulsa, OK, USA). Usually nonparametric tests (Wilcoxon test for two paired groups, Mann-Whitney test for two unpaired groups, Spearman correlation) were used, which do not make assumptions about the distribution of data. For data with mixed-factor design, two-way repeated measures ANOVA test was used with Tukey’s test for post hoc comparisons; in these analyses all data passed the Levene test. All statistical tests were two-tailed. Differences were considered significant when p<0.05. In all figures, population data are represented by symbols and error bars showing mean ± SEM *: p<0.05; **: p<0.01; ***: p<0.001.

## Acknowledgements

We thank Z. Varga-Németh for excellent technical assistance, and B. B. Ujfalussy and Z. Nusser for helpful discussions and comments on the manuscript. This work was supported by the European Research Council (ERC) under the European Union’s Horizon 2020 research and innovation programme (CoG, grant agreement No 771849 to J.K.M.), the International Research Scholar Program of the Howard Hughes Medical Institute (55008740 to J.K.M.) and the NKFIH (K-124824 to J.K.M.).

## Author contribution

A.M., N.K. and J.K.M. performed experiments; B.L. developed analysis procedures. A.M. and J.K.M. analyzed electrophysiology and imaging data. J.K.M. conceived and supervised the project. A.M. and J.K.M. wrote the paper with input from all authors.

## Data availability

Source data are provided for all relevant figures.

## Figure supplements

**Fig. 1 – figure supplement 1.**
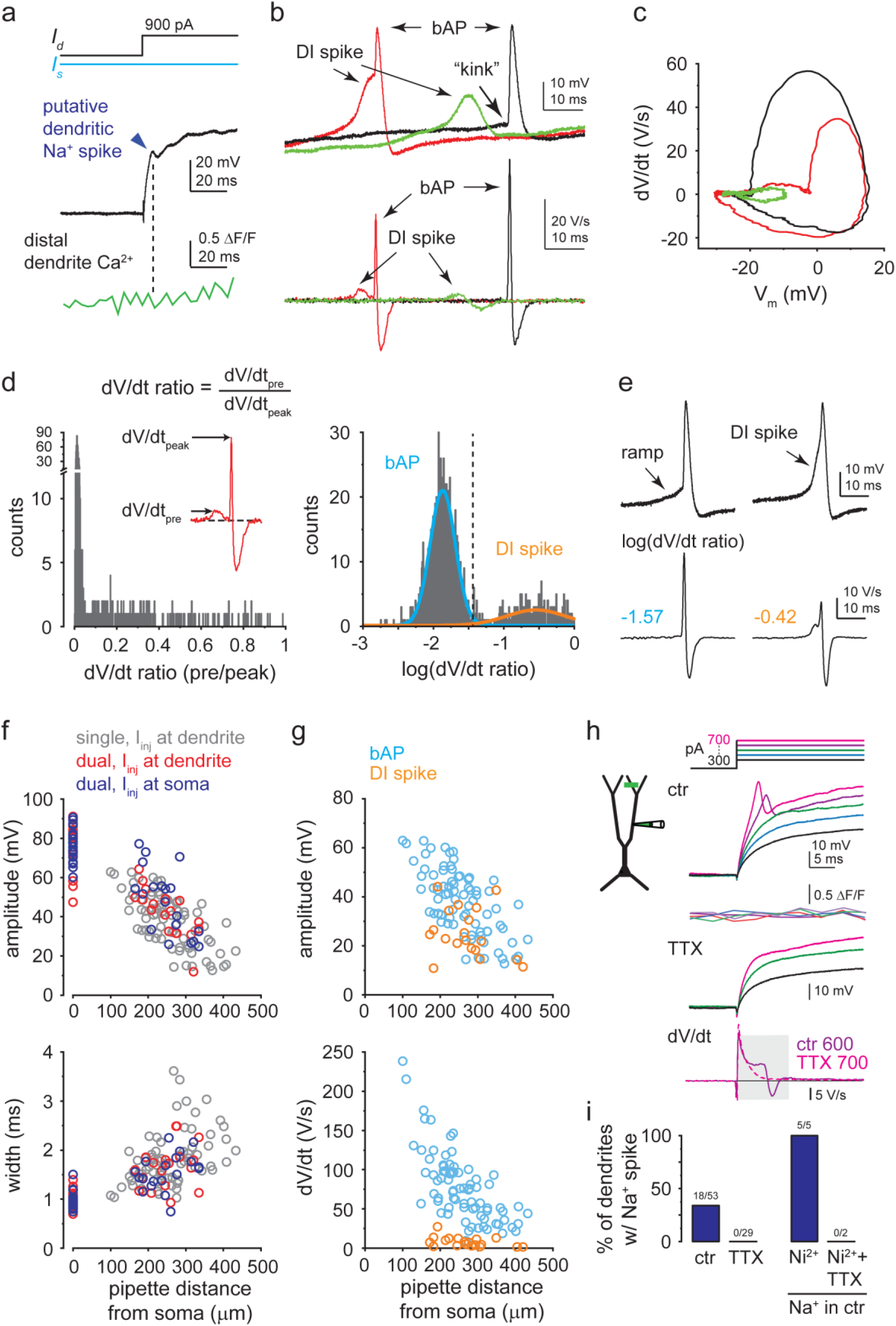
Characteristics of different regenerative dendritic events. **a)** Representative trace from a dual recording, showing the lack of distal dendritic Ca^2+^ signal associated with a putative dendritic Na^+^ spike (dark blue arrow). **b)** Voltage and corresponding dV/dt traces (all recorded from the same CA3PC dendrite), showing a simple bAP (black trace), an isolated DI Ca^2+^ spike (green trace) and a DI Ca^2+^ spike evoking a bAP (red trace). **c)** Phase plot of the traces shown in **b**. Note that the rise of the DI Ca^2+^ spike is slower and more gradual than that of the bAP. **d)** Left, histogram of the dV/dt ratio (defined for each regenerative event as the ratio of the maximum dV/dt value in the 1.5-9 ms time window preceding the peak, to the peak dV/dt). The histogram contains 1206 regenerative events recorded in 16 CA3PCs (between 48-116 events in each cell) expressing both DI spikes and bAPs. Right, After logarithmic transformation of the dV/dt ratio, the histogram can be well fitted by a combination of two Gaussian distributions that belong to simple bAPs (blue) and DI spikes (orange, pooled with or without evoked bAP). Events with log(dV/dt ratio) > −1.44 (dashed line; a threshold corresponding to the mean + 2 s.d. of the bAP peak), were considered to be DI spikes. Categorization by this simple criterion well matched with subjective categorization of events by visual inspection (<0.2% mismatch of events that were confidently categorized to either group by human eye). False positives were due to random fast V_m_ fluctuations in the time window preceding the bAP, whereas false negatives were caused by unusually slow rise kinetics of DI spikes in some cases. **e)** In ∼2% of events a bAP was preceded by a slow but steady ramp, resembling a starting slow dendritic Ca^2+^ spike. If the dV/dt ratio of these events remained <0.035, they were not considered as DI spikes. **f)** Distance dependence of the amplitude (top) and width (bottom) of bAPs. BAPs were initiated by I_inj_ either at the soma (red: dual recordings) or at the dendrite (blue: dual recordings; gray: dendrite-only recordings in an extended dataset). **g)** Comparison of amplitude (top) and dV/dt (bottom) of bAPs and DI spikes (dendrite-only recordings). **h)** Dendritic Na^+^ spike in a representative dendrite-only recording. Increasing I_inj_ steps (protocol on top) produce fast short-latency spikes that are not accompanied by distal dendritic Ca^2+^ signals and that are eliminated by 1 μM TTX. Na^+^ spikes were identified by their clear peak followed by decreasing voltage (negative dV/dt) within the first 10 ms (bottom, averaged traces). **i)** Left, percent of dendrites expressing local Na^+^ spikes to 600-800 pA I_inj_ under control conditions and after addition of TTX. Right, Na^+^ spikes were retained after application of 200 μM Ni^2+^ (n=5 dendrites with Na^+^ spike in ctr), but were eliminated by subsequent application of TTX (n=2).

**Fig. 3 – figure supplement 1.**
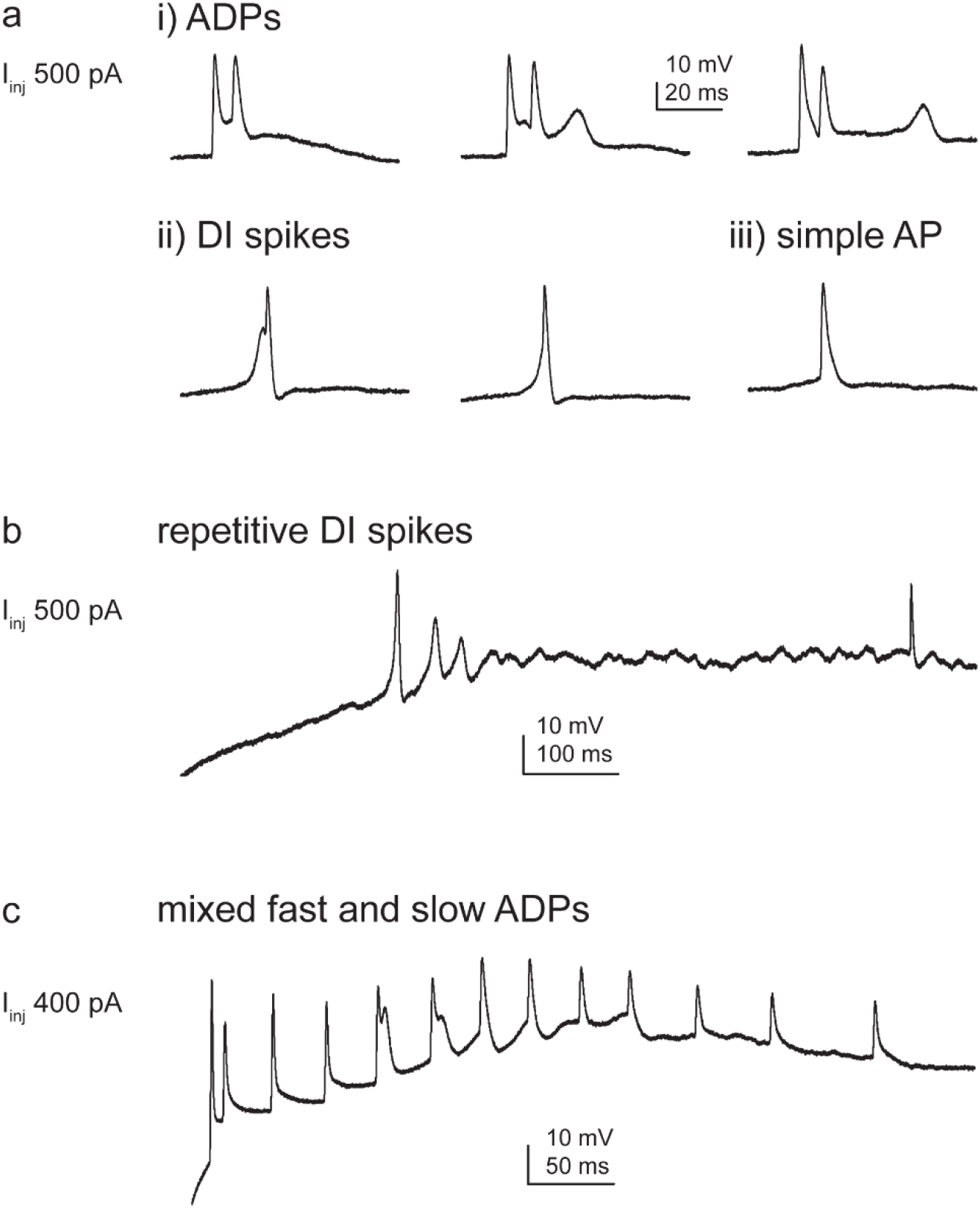
Variability of Ca^2+^ spike phenotypes across and within CA3PCs. **a-c)** Additional examples of CA3PCs expressing variable forms of dendritic Ca^2+^ spikes. **a)** A dendrite expressing both (i) ADPs with slow and fast kinetics, as well as (ii) DI spikes. A simple AP is also shown (iii). All responses were evoked with 500 pA I_inj_. **b)** Some cells fired repetitive DI spikes with progressively smaller amplitude and slower kinetics, eventually resulting in a continuous “noisy” steady-state. **c)** Another dendrite where both fast ADPs and a slowly developing depolarization were induced by I_inj_.

**Fig. 3 – figure supplement 2.**
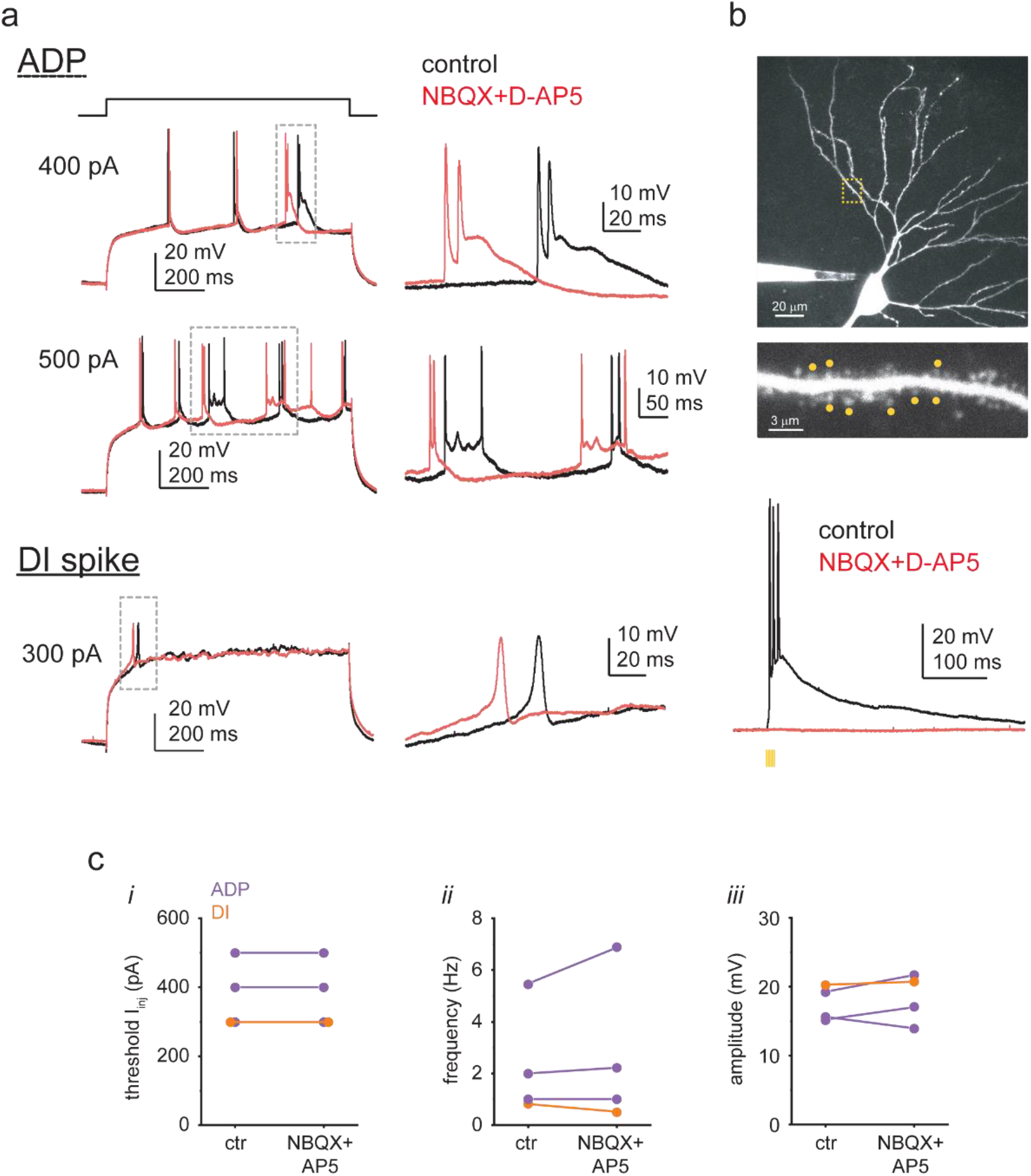
Dendritic Ca^2+^ spike properties do not depend on excitatory synaptic activity. **a)** Representative voltage responses to I_inj_ in a CA3PC dendrite with ADP (top) and another CA3PC dendrite with DI spike (bottom) under control conditions (black) and after bath application of 10 μM NBQX and 50 μM D-AP5 (red; same I_inj_ levels). Dashed boxes are shown at a larger scale on the right. **b)** The receptor blockers eliminated 2PGU-evoked responses as a positive control. Top: z-stack of a somatically recorded CA3PC, and basal dendritic segment (from yellow dashed box) showing the 8 spines stimulated by 2PGU (yellow dots). Bottom: quasi-synchronous uncaging (0.3 ms uncaging, 0.1 ms galvo movement between spines) at the 8 spines evoked large depolarization under control conditions (black) but no response after bath application of 10 μM NBQX and 50 μM D-AP5 (red; same laser power). **c)** Summary of the effect of NBXQ+D-AP5 on (*i*) the threshold I_inj_ level evoking Ca^2+^ spikes, and (*ii*) spike frequency and (*iii*) spike amplitude at threshold I_inj_.

**Fig. 3 – figure supplement 3.**
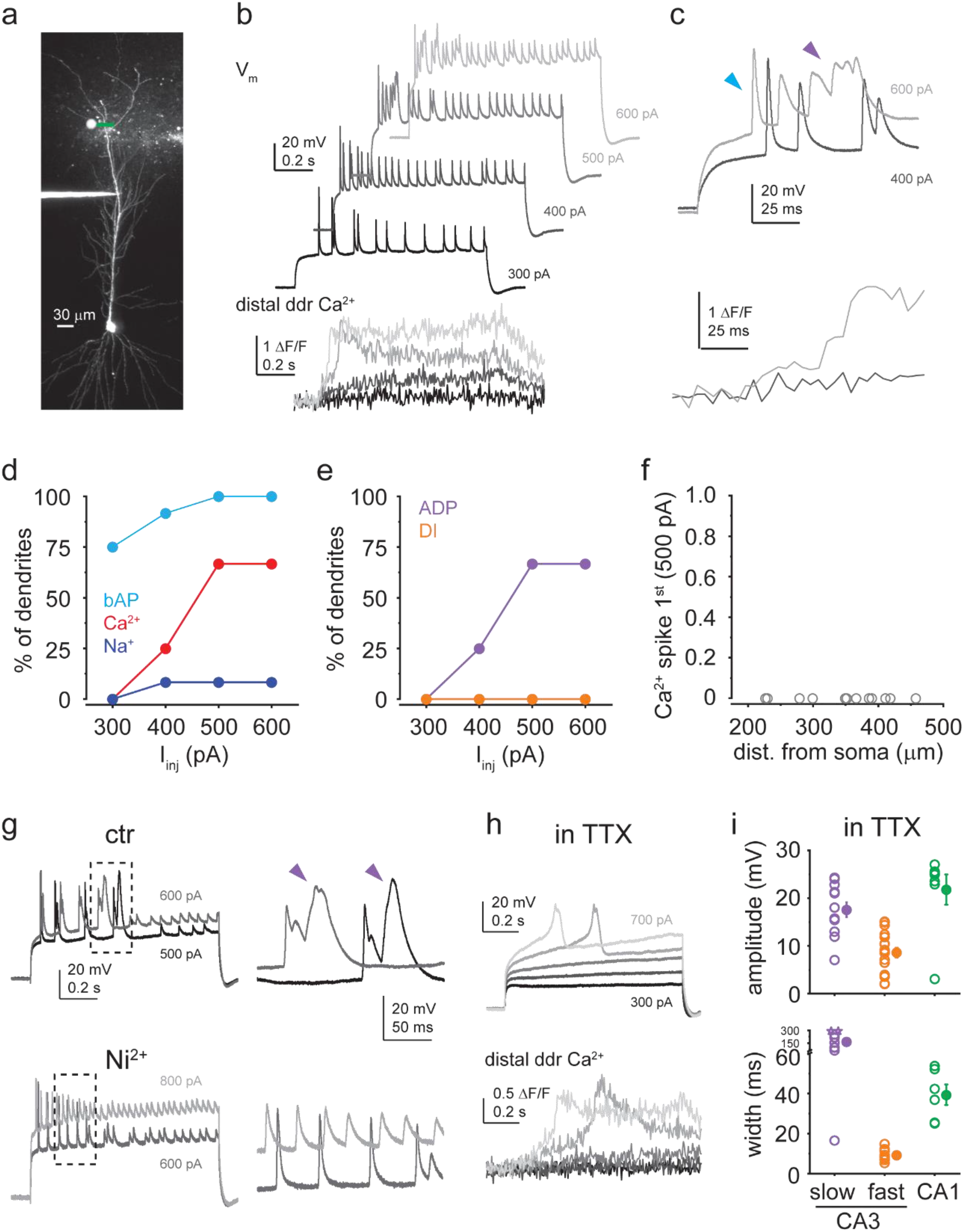
Dendritic Ca^2+^ spikes in CA1PCs are ADP-type. **a)** 2P stack of a CA1PC loaded with OGB-1 and Alexa 594 via a dendritic patch pipette. **b)** Dendritic V_m_ (upper) and distal dendritic Ca^2+^ (lower) traces measured in the cell shown in **a** in response to 1-sec-long step I_inj_ (300-600 pA as indicated). **c)** First parts of the responses to the 400 pA and 600 pA steps are shown enlarged. Note the prolonged depolarization associated with dendritic Ca^2+^ signal upon ADP-like Ca^2+^ spike activation. Arrowheads indicate the first bAP (blue) and the ADP-type Ca^2+^ spike (purple). **d-f)** Summary data from n=12 dendrites. **d)** Percent of dendrites exhibiting bAP (light blue), dendritic Ca^2+^ spike (red) and dendritic Na^+^ spike (dark blue). **e)** Percent of dendrites with ADP-type (purple) and DI (orange) Ca^2+^ spikes. **f)** Probability of Ca^2+^ spikes being the first regenerative event evoked by 500 pA I_inj_ steps. No DI Ca^2+^ spikes were detected. **g)** Left, V_m_ responses to two I_inj_ steps under control conditions (upper) and after bath application of 200 μM Ni^2+^ (lower), Right: Enlarged regions in the boxes on the left panel. Note that the ADP-type Ca^2+^ spikes (indicated by purple arrows) observed in the control solution were inhibited by Ni^2+^ (n=3 experiments). **h)** Ca^2+^ spikes measured in 1 μM TTX in the bath. I_inj_ was increased in 100 pA steps from 300 to 700 pA. Note the dendritic Ca^2+^ signals associated with the Ca^2+^ spikes. **i)** Comparison of dendritic Ca^2+^ spike amplitude and width (as measured in TTX) in CA1PCs (green, n=7/6 for ampl./width) versus slow (purple, n=12) and fast (orange, n=18/13 for ampl./width) Ca^2+^ spikes in CA3PCs. Width was measured on spikes with >5mV amplitude. Note that CA1PC Ca^2+^ spikes have similar amplitude but intermediate duration between fast and slow CA3PC spikes. Data in a-c, g and h are from three different cells.

**Fig. 5 – figure supplement 1.**
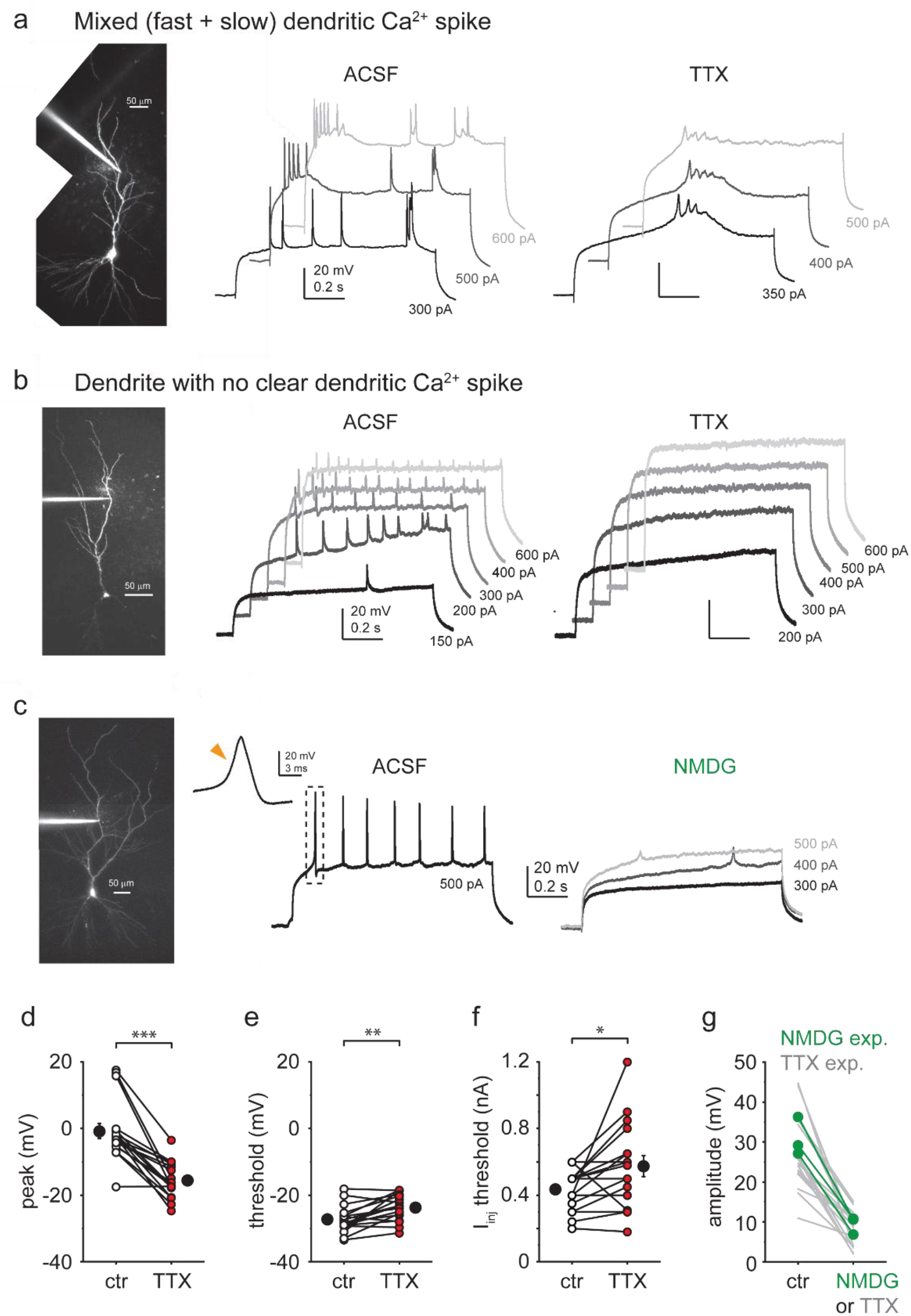
Additional properties of dendritic Ca^2+^ spikes. **a)** Example recording of a CA3PC dendrite with mixed types of Ca^2+^ spikes in TTX. **b)** Example recording of a CA3PC dendrite with no clear Ca^2+^ spike. **c)** Example recording of a CA3PC dendrite with DI spike in ACSF and after partial replacement of Na^+^ with NMDG^+^. **d-f)** Effect of TTX on peak (**d**; mean difference ∼15 mV, n=17), voltage threshold (**e**; mean difference ∼3.5 mV, n=17) and I_inj_ threshold (**f**, mean difference: ∼140 pA, n=17) of DI fast spikes. **g)** Summary of the impact of NMDG^+^ on DI spike amplitude (n=3). Note that the effect is similar to that of TTX (data from Fig. 5d).

**Fig. 8 – figure supplement 1.**
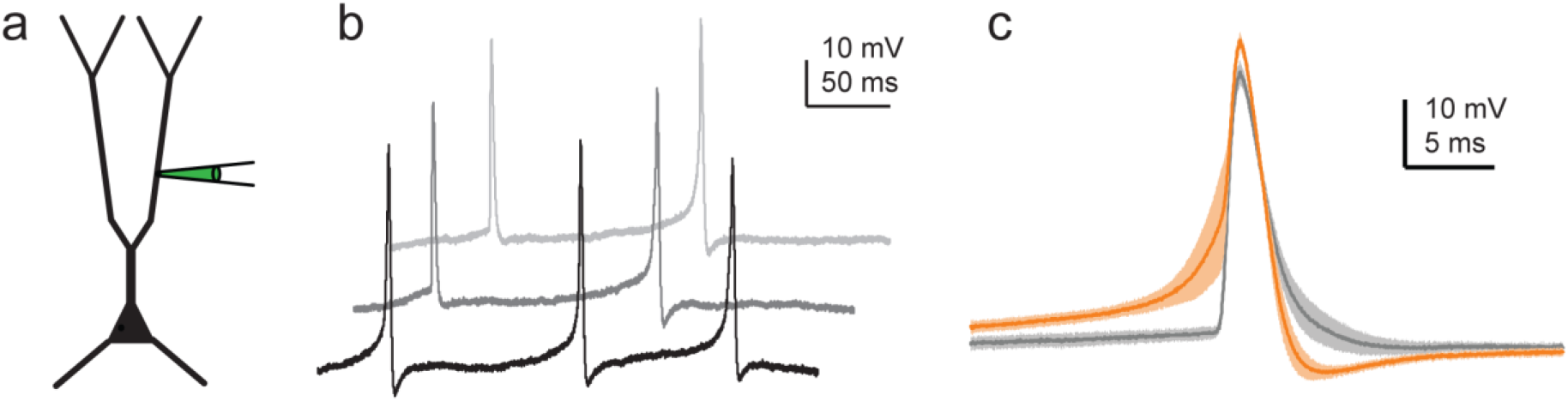
Effect of DI spikes on AP output in single-site dendritic recordings. **a)** Experiment schematic. **b)** Three individual trace segments during 200 pA I_inj_ steps recorded in a representative CA3PC dendrite, showing s-APs and Ca-APs. **c)** Ca-APs (orange, mean ± s.e.m. of n=15 events) and s-APs (grey, mean ± s.e.m. of n=10 events) aligned to peak from the experiment shown in b.

